# A divergent *Plasmodium* NEK4 acts as a key regulator driving the early events of meiosis

**DOI:** 10.1101/2025.11.21.689802

**Authors:** Ryuji Yanase, Molly Hair, Mohammad Zeeshan, David J. P. Ferguson, Declan Brady, Carla Pasquarello, Andrew Bottrill, Suhani Bhanvadia, Arrmund Neal, Eelco C. Tromer, Karine G. Le Roch, Alexandre Hainard, Anthony A. Holder, Sue Vaughan, David S. Guttery, Rita Tewari

**Affiliations:** School of Life Sciences, Queen’s Medical Centre, University of Nottingham, Nottingham, UK; Department of Genetics, Genomics and Cancer Sciences, College of Life Sciences, University of Leicester, Leicester, UK; Oxford Brookes University, Department of Biological and Medical Sciences, Oxford, UK; Proteomics Core Facility, Faculty of Medicine, University of Geneva, Switzerland; School of Life Sciences, Gibbet Hill Campus, University of Warwick, Coventry, UK; Department of Molecular, Cell and Systems Biology, University of California, Riverside, Riverside, United States; Cell Biochemistry, Groningen Institute of Biomolecular Sciences & Biotechnology, University of Groningen, Groningen, The Netherlands; Malaria Parasitology Laboratory, The Francis Crick Institute, London, UK

**Author notes:** Corresponding authors (RT); (DSG).

**Keywords:** *Plasmodium*, meiosis, NIMA-related kinase, malaria, zygote, ookinete, MTOC, microtubules, chromosome

## Abstract

Meiosis is a conserved yet evolutionarily varied process underpinning sexual reproduction in eukaryotes. In the malaria parasite *Plasmodium*, meiosis is unconventional: it occurs immediately after fertilisation (post-zygotic) and must be coordinated with the transformation of the zygote into a motile ookinete. The mechanisms synchronising these meiotic and morphogenetic programmes remain unknown. Here, we identify the *Plasmodium berghei* NIMA-related kinase NEK4 as a key regulator that couples meiotic initiation with zygote morphogenesis. Using ultrastructure expansion microscopy, we show that NEK4 accumulates at the microtubule-organising centre (MTOC) and the apical polar complex (APC) shortly after fertilisation, preceding the assembly of perinuclear and cortical microtubules. We reveal that *Plasmodium* zygotes undergo MTOC-associated nuclear migration, analogous to the meiotic nuclear movement in fission yeast. Deletion of the *Pbnek4* gene results in complete developmental arrest: MTOC duplication and microtubule formation are blocked, chromatin remains uncondensed, and nuclear migration and cell polarity fail to establish. Transcriptomic and phosphoproteomic analyses reveal that absence of NEK4 causes a collapse in transcriptional and phosphoregulatory networks governing meiosis and cytoskeletal organisation, leading to reduced expression and phosphorylation of important players, including HOP1, REC8, and AP2-O. These findings establish NEK4 as a key regulator driving meiotic entry and zygote maturation.

## Introduction

Meiosis is a conserved and essential mode of cell division that underpins sexual reproduction in eukaryotes. It entails a carefully choreographed sequence of events, including DNA replication, homologous recombination, pairing of homologous chromosomes (synapsis) and two rounds of chromosome segregation^1^. In mammals, it generates haploid gametes from diploid progenitors, following two successive nuclear divisions without intervening genome duplication^1^. In many well-studied organisms including yeast, plants, and mammals, meiotic progression is regulated by a network of conserved protein kinases and checkpoints, including cyclin-dependent kinases (CDKs), Polo-like kinases (PLKs), the anaphase-promoting complex/cyclosome (APC/C), ataxia-telangiectasia mutated (ATM)/ATM and Rad3-related (ATR) kinases, checkpoint kinases (CHK1/2), and members of the NIMA-related kinase (NEK) family^2, 3, 4, 5^. These regulators ensure the fidelity of chromosome segregation by integrating signals from DNA damage checkpoints, spindle dynamics, and crossover/recombination status, at the same time as maintaining synchrony with cytoskeletal changes that support cell division and differentiation.

Despite the broad conservation of its core function to facilitate obligatory chromosomal exchanges and homologous recombination between parental chromosomes, both the molecular machinery executing meiosis and its operational sequence of events display remarkable variability^6, 7^. For a wide range of unicellular eukaryotes, including parasitic protists, meiotic programmes have undergone substantial evolutionary divergence. This is evident in the malaria-causing parasite *Plasmodium*, a member of the phylum Apicomplexa^7,8^. This haploid parasite has a complex lifecycle with proliferative asexual multiplication in the liver and blood stream of the host^9^. A subpopulation of blood-stage parasites differentiates into haploid male and female gametocytes which, following ingestion in a mosquito blood meal, start the sexual phase of the life cycle inside the gut of the vector. Gametogenesis produces haploid gametes and following fertilisation, meiosis occurs in the diploid zygote; a process known as post-zygotic meiosis. The zygote undergoes a ∼24-hour developmental programme to become an elongated, motile ookinete capable of traversing the midgut epithelium^8^ (Fig. 1a), where it forms an oocyst. Many haploid sporozoites then emerge from the oocyst, migrate to the mosquito salivary glands and transmit to the vertebrate host in the blood meal. *Plasmodium* infections result in over 250 million clinical cases and nearly 600,000 deaths each year, disproportionately affecting young children in sub-Saharan Africa^10^. These outcomes are driven not only by the rapid asexual proliferation in the human host, but also by the successful sexual reproduction and transmission through the mosquito vector. Transmission is an obligate part of the life cycle and therefore prevalence of the disease is critically reliant on the process of meiosis^9^.

**Figure 1.**
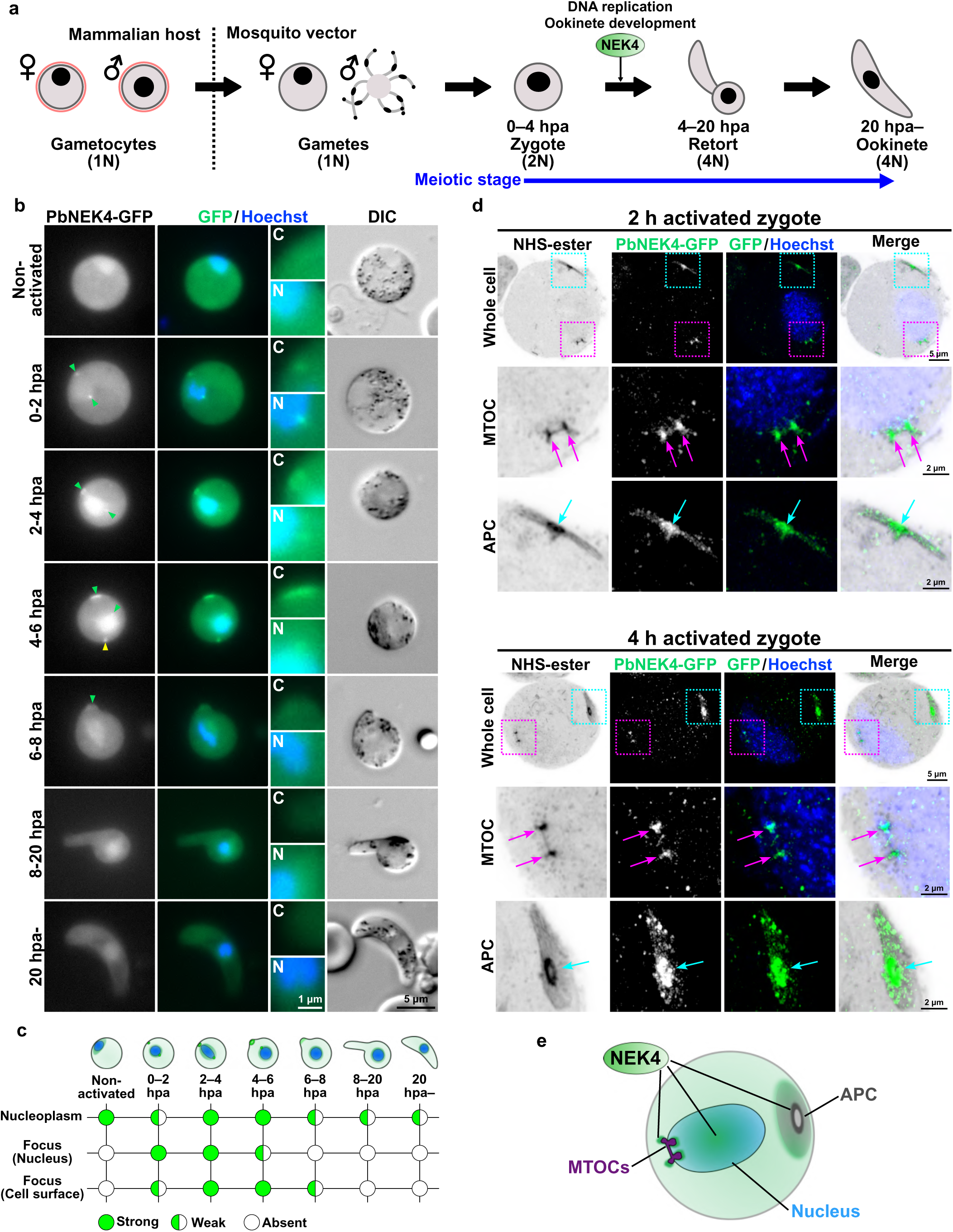
NEK4 is located at the MTOC, APC, and nucleus during the early meiotic phase of zygote to ookinete development. **a.** A schematic diagram of ookinete development showing formation of gametocytes, gametes, and zygote, and the involvement of NEK4 in cellular development and meiosis. The developmental stage and nuclear ploidy (from 1N to 4N) are indicated for each phase. Activation of gametocytes in the mosquito gut initiates sexual development. In the schematic, the red blood cell membrane is depicted in red, the parasite cytoplasm in grey, and the nucleus as a black circle. The time scale in hours post-activation (hpa) is also indicated for the developmental stages from zygote to retort and ookinete. **b.** Live-cell imaging of PbNEK4-GFP location at different time points (hpa). Green arrowheads indicate the PbNEK4-GFP foci in the nucleus (Hoechst, blue) and at the cell periphery. Insets show magnification of the cell periphery (C) and nucleus (N). The yellow arrowhead indicates the cytoplasmic foci of PbNEK4-GFP. Images are representative of 30–50 cells analysed across at least 3 independent biological experiments. **c.** Schematic summary of the location of PbNEK4-GFP shown in (b), indicating the signal strength in the nucleoplasm, nuclear focus, and at the cell periphery focus over time. **d.** Expansion microscopy of PbNEK4-GFP zygotes at 2 hpa and 4 hpa. The cells were labelled with NHS-ester (gray), anti-GFP antibody (green), and Hoechst (blue). The whole-cell images display the MTOC and APC regions enclosed by superimposed magenta and cyan dashed boxes, respectively. The corresponding magnified views of these regions are shown below. The positions of the MTOC and APC are indicated by magenta and cyan arrows, respectively. Representative images from three independent biological experiments (n ≥ 5 cells examined per condition). **e.** Model illustrating the localisation of NEK4 at the MTOCs, APC and nucleoplasm.

Meiosis in *Plasmodium* is divergent from the canonical process observed in model eukaryotes. In these organisms, a diploid cell undergoes pre-meiotic DNA replication followed by prophase I substages including leptotene, zygotene, pachytene, diplotene and diakinesis, before metaphase I and anaphase I of the first meiotic division and then meiosis II, ultimately generating four separate germ cells, each containing a haploid nucleus^11^. In *Plasmodium*, eukaryote mostly haploid and meiosis begins in diploid zygote formed after fertilisation. The early meiotic events, for example DNA replication, chromosome pairing, and formation of chromosome axes (proteinaceous scaffolds that form along sister chromatids), are initiated within the first few hours post-fertilisation. Ultrastructural studies have shown that *Plasmodium* zygotes progress through early prophase I stages (leptotene and zygotene), forming distinct synaptonemal complexes (SC) that persist until metaphase I^12, 13^, whereas in canonical meiosis the SC assembles during zygotene, is fully established at pachytene, and normally disassembles during diplotene before diakinesis and metaphase I^11^. However, in *Plasmodium* later prophase I stages (including diplotene and diakinesis) and canonical meiosis II have not been observed^14, 15^. Despite two rounds of chromosome segregation, the ookinete retains a single nucleus with four kinetochore clusters^16, 17^, indicating a divergent closed meiosis without karyokinesis (similar to meiosis I in *S. pombe* and the closed mitosis characteristic of *Plasmodium*^17^), rather than the formation of four separate haploid nuclei.

The cell biological divergence of meiosis is reflected at the molecular level as well. *Plasmodium* species lack orthologues of several conserved meiotic regulators, including CDC25, CDC14, PLKs, and many APC/C components^18, 19^. They do possess a reduced, parasite-specific set of protein kinases that appear to have acquired regulatory roles normally fulfilled by these missing elements^19, 20^. The signalling mechanisms that control *Plasmodium* meiosis and coordinate with cytoskeletal reorganisation and establishment of cell polarity, are poorly defined, although cGMP signalling has been suggested to initiate ookinete gliding motility and polarization^21^. One protein kinase family of particular interest is the NIMA-related kinases (NEKs), which are conserved across eukaryotes and regulate centrosome function, spindle assembly, and 3D chromatin organisation^22, 23^. In metazoans, specific NEKs such as NEK1, NEK2, and NEK9 function in meiosis and mitosis, often through regulation of microtubule dynamics and DNA damage (NEK1) and spindle assembly (NEK2 and NEK9) checkpoint control^22, 24^. The *Plasmodium* genome encodes four NEKs (NEK1–4), which have, apart from *Plasmodium* NEK1 and human NEK2, no direct 1-to-1 human orthologs (Extended Data Fig. 1a, b). Each *Plasmodium* NEK has a distinct expression pattern and function during the parasite life cycle^25, 26, 27, 28^, with NEK2 and NEK4 shown to be critical for zygote development and parasite transmission^19, 25^ (Extended Data Fig. 1c). Deletion of the *Pbnek4* gene results in a failure of DNA replication in the zygote following fertilisation^25^ (Figure 1a), yet the underlying function and phosphosignalling network remain undefined.

Given its post-zygotic meiosis within a haploid life cycle and the lack of several canonical meiotic regulators^8^, Plasmodium provides a powerful model to investigate how minimal signalling networks coordinate chromosome dynamics with cytoskeletal reorganisation. Here, we show that during early zygote development where the post-zygotic meiosis commences, *Plasmodium berghei* (Pb) NEK4 is located at the microtubule organising centre (MTOC) and apical polar complex (APC). Deletion of the *Pbnek4* gene disrupts microtubule organisation, MTOC development, and nuclear migration during meiotic S phase and prophase. Furthermore, its absence impairs transcription and phosphorylation of key meiotic proteins, leading to a failure in chromosome condensation and developmental arrest at the earliest stages of meiosis. We propose that this kinase integrates the nuclear and cell differentiation events that drive meiotic progression and ookinete development, respectively.

## Results

### NEK4 moves to the nucleus, MTOC, and apical polar complex during early meiotic development

Our previous study showed that NEK4 ablation blocks zygote development^25^. To investigate NEK4 functions during the meiotic stages of *P. berghei* development, first we determined the spatiotemporal location of NEK4, using a transgenic parasite line expressing the protein with a C-terminal GFP tag from the modified endogenous gene locus (PbNEK4-GFP – Extended Data Fig. 1d-f). Live-cell imaging revealed a dynamic pattern of PbNEK4-GFP location following female gametocyte activation, zygote differentiation and ookinete development (Fig. 1b and Extended Data Fig. 2 and 3a). In non-activated female gametocytes, the GFP signal was diffuse throughout the nucleoplasm and cytoplasm (Fig. 1b). Then shortly after fertilisation and in the first two hours post-activation (hpa), the PbNEK4-GFP signal was concentrated in two foci (Fig. 1b), one at the periphery of the nucleus near the nuclear envelope and the other at the cell cortex. By two to six hpa, these foci were more pronounced, suggesting a localisation of PbNEK4-GFP at both nuclear and apical regions of the cell during this period of intense morphological transformation. The PbNEK4-GFP signal within the nucleoplasm appeared stronger relative to the cytoplasm. In addition, a focal dot-like PbNEK4-GFP-positive structure was observed moving within the cytoplasm (Fig. 1b; yellow arrowhead and Supplementary Video 1 and Extended Data Fig. 3b). This structure remained visible during the early stages of zygote-ookinete development (until ∼10 hpa) but was no longer detectable in the fully mature ookinete (Extended Data Fig. 2 and 3c). We speculate that this focus may represent a remnant of the NEK4-associated structure that undergoes degradation during the stages of development. The morphological transformation of the zygote into an ookinete begins at about 4 hpa and is initiated at the site where PbNEK4-GFP is located at the cell periphery. The onset of this transformation is accompanied by reduction of this GFP signal and the disappearance of the nuclear PbNEK4-GFP focus, with residual PbNEK4-GFP dispersed diffusely throughout the cytoplasm and nucleoplasm (Fig. 1b, c, and Extended Data Fig. 3a).

To examine in more detail the location of NEK4 within the developing zygote, we used ultrastructure expansion microscopy (U-ExM). We utilised NHS-ester staining, which labels primary amine groups on proteins to reveal the global intracellular architecture like electron microscopy^29^. This staining clearly revealed the nuclear MTOC and the development of the APC at the cell periphery. Immunofluorescence labelling of the U-ExM samples with an anti-GFP antibody confirmed that PbNEK4-GFP was located at both the MTOC and the APC (Fig. 1d). Furthermore, at 4 hpa, the dividing and separating MTOCs and a more developed APC were observed by NHS-ester staining. At this later time point, a distinct presence of PbNEK4-GFP at both the MTOCs and the APC was confirmed (Fig. 1d). These observations indicate prominent localisation of NEK4 during the early stages of zygote development (2 to 4 hpa), with a concentration at the MTOC and APC, alongside a diffuse distribution within the nucleoplasm (Fig. 1e).

### NEK4 coordinates nuclear repositioning and directional microtubule formation in the zygote

In our study of NEK4 localisation in early meiotic zygotes, we observed that the nucleus shows dynamic movement during early meiosis as seen across various eukaryotes^6, 30, 31^. For example, in the fission yeast *Schizosaccharomyces pombe* (*S. pombe*), nuclear movement during meiosis is characterised by a dramatic “horsetail” motion, where the nucleus oscillates back and forth along the cell axis during prophase I^32, 33^. This movement is driven by cytoplasmic dynein and microtubules anchored at the spindle pole body (SPB)^34, 35^, and is essential for efficient homologous chromosome pairing and recombination^32^. Time-lapse imaging of PbNEK4-GFP zygotes at 2 hpa revealed that inside the zygote, PbNEK4-GFP focus localised at the nucleus and moves together with the nucleus (magenta arrowheads in Fig. 2a and Supplementary Video 2, 3). To examine MTOC and microtubule dynamics in parallel during the nuclear movement, we used a transgenic *P. berghei* line expressing EB1 (End Binding protein 1) fused to GFP (PbEB1-GFP)^16^, EB1 is a conserved microtubule plus-end binding protein, but in *Plasmodium*, EB1 marks the microtubule lattice and the MTOC^16, 36^. The PbEB1-GFP focus, representing the MTOC position, was observed moving within the zygote, leading the nucleus (magenta arrowheads in Fig. 2b). The nucleus appeared elongated, seemingly pulled by the PbEB1-GFP focus. Additionally, PbEB1-GFP signal was observed extending from the focus, likely representing microtubules extending from the MTOC (Fig. 2b and Supplementary Video 4, 5). Furthermore, the fact that the localisation of PbNEK4-GFP and PbEB1-GFP at the cell periphery remains largely stationary contrasts with the nuclear movement (cyan arrowheads in Fig. 2a and b), indicating that this nuclear movement is independent of whole-cell rotation or migration.

**Figure 2.**
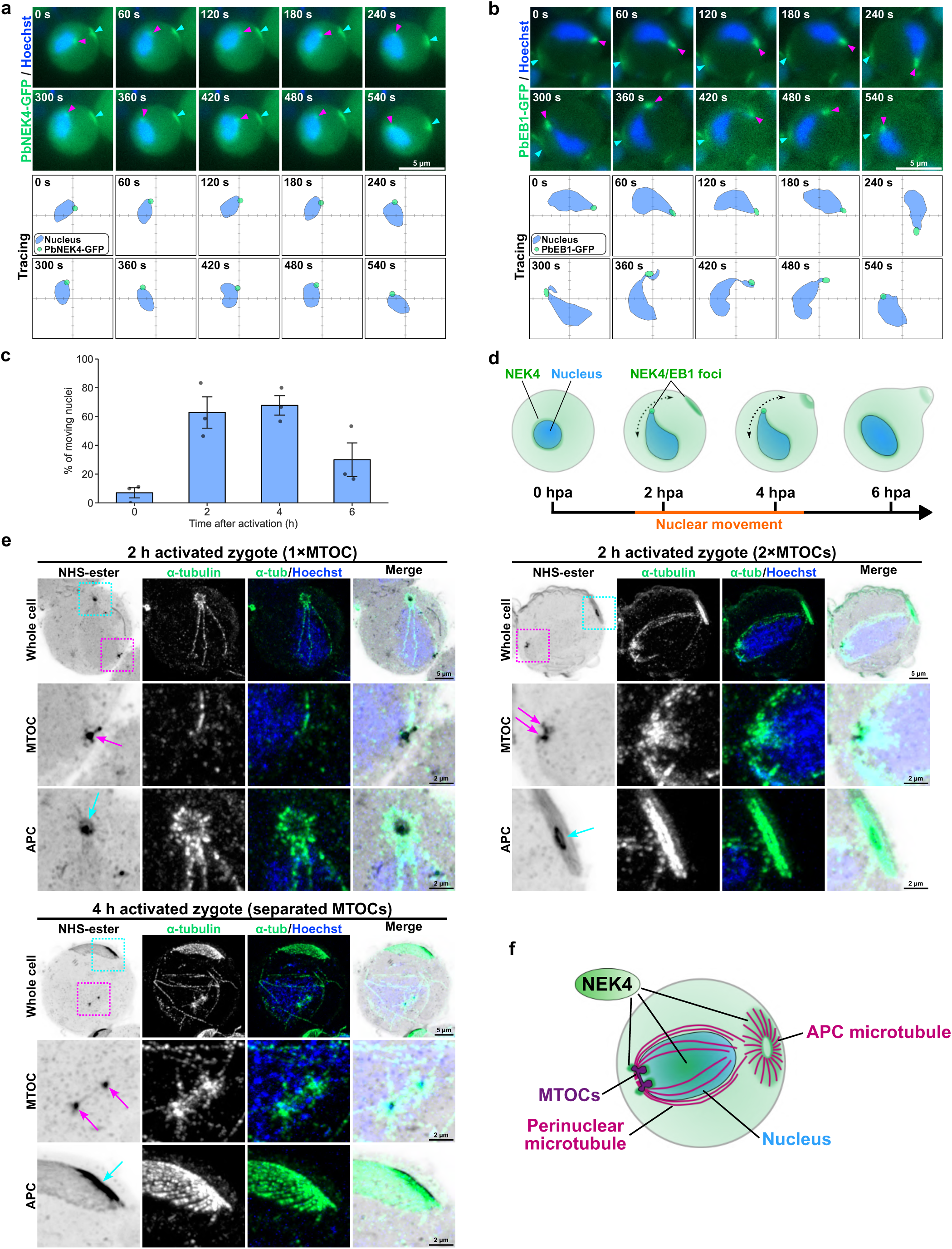
NEK4-localised MTOC leads nuclear movement, and directional microtubule formation occurs at both the MTOC and the APC during early meiosis. **a.** Time-lapse imaging of a PbNEK4-GFP expressing zygote shows nuclear movement. The PbNEK4-GFP foci in the nucleus and at the cell periphery are indicated by magenta and cyan arrowheads, respectively. The lower panel (“Tracing”) tracks the position of the PbNEK4-GFP focus (green dot) and the nucleus (blue) over 540 seconds. **b.** Time-lapse imaging of a zygote expressing the microtubule plus-end/MTOC marker PbEB1-GFP (green). The PbEB1-GFP foci in the nucleus and at the cell periphery are indicated by magenta and cyan arrowheads, respectively. The lower panel (“Tracing”) tracks the position of the PbEB1-GFP focus (green dot) and the nucleus (blue) over 540 seconds. **c.** Quantification of the percentage of zygotes showing nuclear movement at 0, 2, 4, and 6 hpa. The data represent the mean ± SEM from three independent experiments (n = 3), in which 30 cells were analysed per experiment. Black dots show the result from each experiment. **d.** Schematic model of nuclear movement, illustrating the association of NEK4/EB1 foci with the moving nucleus at 2-4 hpa. **e.** Expansion microscopy showing microtubule organisation in developing zygotes. Microtubules (α-tubulin, green) are shown at 2 hpa with either one or two MTOCs, and at 4 hpa with separated MTOCs. The whole-cell images display the MTOC and APC regions enclosed by superimposed magenta and cyan dashed boxes, respectively. The corresponding magnified views of these regions are shown below. Representative images from three independent biological experiments (n ≥ 5 cells examined per condition). The positions of the MTOC and APC are indicated by magenta and cyan arrows, respectively. **f.** Model of the microtubule organisation in a developing zygote, illustrating the formation of perinuclear and APC microtubules and the location of PbNEK4-GFP at the MTOCs, APC and nucleoplasm.

Quantification of the proportion of zygotes at different stages during development with motile nuclei indicated that the movement peaked at between 2 and 4 hpa and then subsided (Fig. 2c). These observations suggest that in *P. berghei* zygotes there is a form of nuclear movement like that widely observed in *S. pombe* and other eukaryotes^6, 30, 31, 32, 37^. This movement appears to be led by the MTOC, where both PbNEK4-GFP and PbEB1-GFP are located (Fig. 2d). We also performed dual-colour live-cell imaging using a parasite line expressing PbNEK4-GFP and PbEB1-mCherry. This analysis revealed a clear colocalization of NEK4 and EB1 signals in the nucleus and at the cell periphery (Extended Data Fig. 4a).

To investigate the microtubule structures at higher resolution during the early development stages, we performed U-ExM using an anti-α-tubulin antibody (Fig. 2e and Extended Data Fig. 4b, c). At 2 hpa, two populations of zygotes were present: those with a single MTOC and those with duplicated MTOCs. In cells with a single MTOC (Fig. 2e, top left panel), multiple microtubules were visible in the perinuclear region, with one microtubule oriented as if extending from the nuclear MTOC toward the APC. In cells with duplicated MTOCs (Fig. 2e, top right panel), both microtubules emanating from the MTOCs and microtubule organisation at the APC appeared more developed. Some of the microtubules running along the nuclear periphery appeared to connect to the APC. In zygotes at 4 hpa (Fig. 2e, bottom left panel), the two MTOCs were separating, with a microtubule structure resembling a spindle oriented between them. Further development of microtubule structures was also evident in the perinuclear region and at the APC. In summary, significant microtubule development occurs at both the nuclear MTOCs and the APC, the sites of greatest PbNEK4-GFP accumulation (Fig. 2f), suggesting that NEK4 may be involved in regulating microtubule polymerisation or dynamics. This association of NEK4 with the MTOC and APC suggests a function as a spatial organiser of microtubule polarity in the developing zygote. The microtubules extending around the nucleus appear analogous to the perinuclear microtubules observed in other eukaryotes during early meiosis, which mediate nuclear and chromosome movement^32, 38^.

### NEK4 is essential for nuclear migration, microtubule formation, and morphological transformation from zygote to ookinete

To assess the importance of NEK4 during zygote/meiotic development, we used an existing *nek4* gene knockout parasite line (*Pbnek4-ko*; Extended Data Fig. 5a, b)^25^. A previous study has shown that while *Pbnek4-ko* parasites undergo fertilisation, subsequent DNA replication and ookinete formation are blocked^25^. Live-cell imaging over the period between 2 to 24 hpa confirms that *Pbnek4-ko* zygotes failed to elongate or transition through the retort form into mature ookinetes, in contrast to PbNEK4-GFP control cells that completed this transformation within 20 hours (Fig. 3a). We then investigated the nuclear movement observed in zygotes. Although moving nuclei were observed in a few *Pbnek4-ko* zygotes between 2 and 4 hpa, the proportion of cells with moving nuclei was significantly reduced compared to the PbNEK4-GFP controls (Fig. 3b).

**Figure 3.**
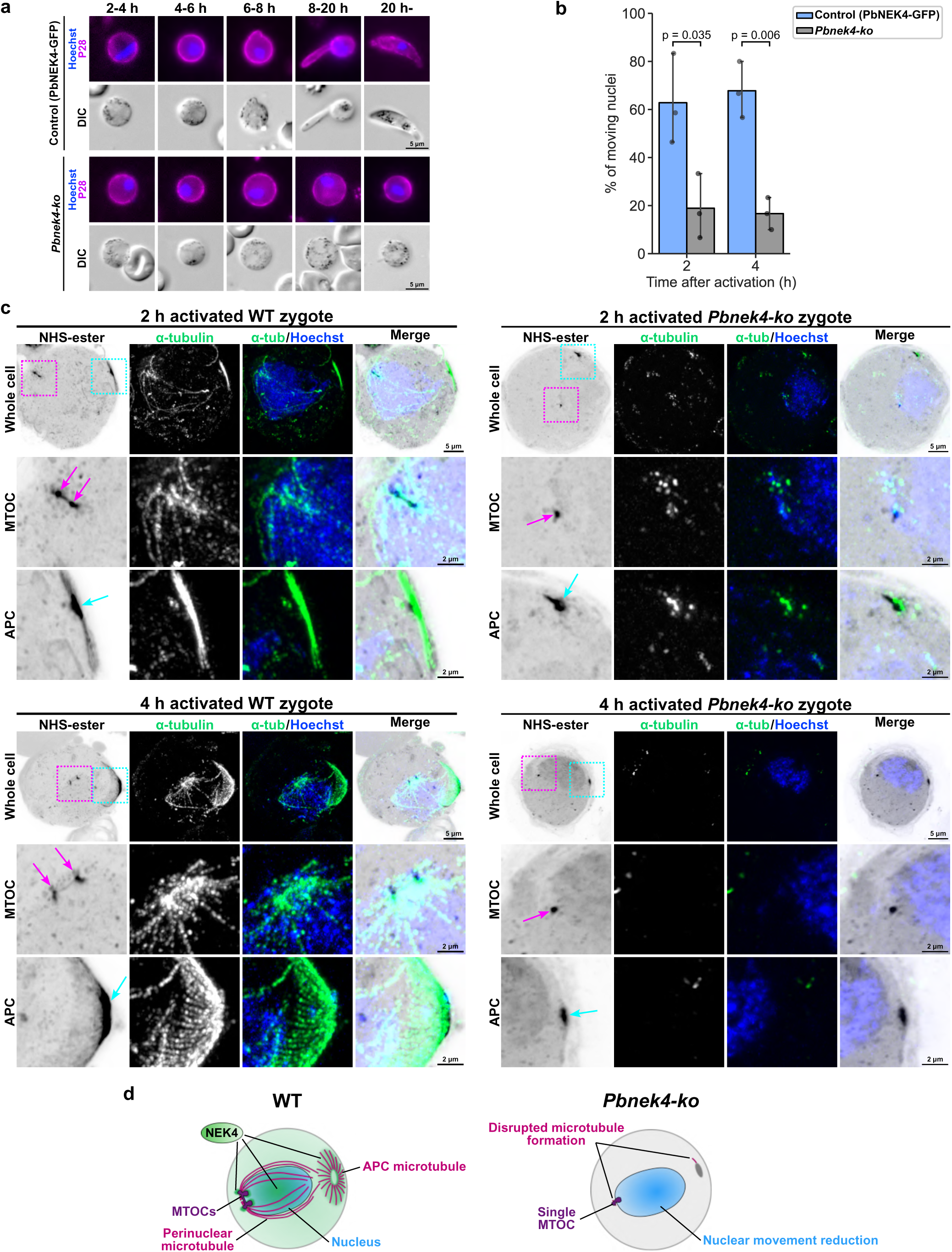
NEK4 is essential for DNA replication, nuclear migration, microtubule formation, and morphological transformation from zygote to ookinete. **a.** Live-cell imaging of control (PbNEK4-GFP) and *Pbnek4-ko* parasites from 2 to 20 hpa. Representative live-cell images of different cells sampled at specific time points are shown. Parasites were labelled with Hoechst (blue) and a Cy3-conjugated 13.1 antibody (magenta), which recognises P28 protein on the surface of zygotes and ookinetes, immediately prior to imaging. Unlike the control which develops into an elongated ookinete, *Pbnek4-ko* parasites remain round and fail to develop. Images are representative of 30–50 cells analysed across at least 3 independent biological experiments. **b.** Quantification of nuclear movement in control (PbNEK4-GFP, blue) and *Pbnek4-ko* (gray) zygotes at 2- and 4 hpa. Nuclear movement is significantly reduced in the *Pbnek4-ko* mutant both at 2- and 4 hpa. The data represent the mean ± SEM from three independent experiments (n = 3), in which 30 cells were analysed per experiment. Black dots show the result from each experiment. P-values were determined by Welch’s t-test. **c.** Expansion microscopy comparing microtubule organisation (α-tubulin, green) in WT and *Pbnek4-ko* zygotes at 2 and 4 hpa. The whole-cell images display the MTOC and APC regions enclosed by superimposed magenta and cyan dashed boxes, respectively. The corresponding magnified views of these regions are shown below. The positions of the MTOC and APC are indicated by magenta and cyan arrows, respectively. *Pbnek4-ko* zygotes show a single MTOC and disrupted microtubule formation, in contrast to the well-organised perinuclear and APC microtubules in WT zygotes. Representative images from three independent biological experiments (n ≥ 5 cells examined per condition). **d.** Schematic model summarising the defects in *Pbnek4-ko* zygotes, including the persistence of a single MTOC, disrupted microtubule formation, and reduced nuclear movement.

To investigate microtubules in *Pbnek4-ko* zygotes, we used U-ExM. This revealed that the distinctive formation of perinuclear and APC-associated microtubules in WT cells was almost completely ablated in *Pbnek4-ko* zygotes (Fig. 3c). Although the presence of some tubulin was detected, the formation of the microtubule-based structures at the APC was almost completely disrupted compared to PbNEK4-GFP controls (WT; Fig. 3c). Transmission electron microscopy (TEM) analysis also confirmed that partial formation of the APC and the presence of microtubules at the APC and the perinuclear region were still observable in *Pbnek4-ko* zygotes (Extended Data Fig. 5c). In non-activated female gametocytes, a single MTOC-like structure was observed in both WT and *Pbnek4-ko* parasites, but neither the formation of microtubules extending from it nor the assembly of the APC structure was evident at this stage (Extended Data Fig. 5d, e). Following activation, *Pbnek4-ko* zygotes exhibited partial formation of the APC and partial assembly of microtubules at the MTOC and APC. Taken together, these results suggest that although *Pbnek4* deletion does not completely block microtubule formation, it profoundly compromises their formation or stability. Additionally, while the formation of a nuclear MTOC was observed, it failed to duplicate and remained as a single MTOC over the first four hours after fertilisation. Together, these data on the consequences of *Pbnek4* gene deletion demonstrate that NEK4 is essential for MTOC duplication and the organisation of perinuclear and apical microtubule networks. These processes are prerequisites for DNA replication, meiotic entry, and morphological transition from zygote to ookinete. In the absence of NEK4, the microtubule-based cytoskeletal framework necessary for meiotic progression fails to assemble, leading to developmental arrest prior to transition to retort form (Fig. 3d).

### NEK4 is essential for establishing the ultrastructure necessary for meiotic chromatin condensation and APC development

To further examine the ultrastructural consequences of *Pbnek4* knockout, we used TEM to study WT and *Pbnek4-ko* zygotes (Fig. 4a). At 4 hpa, WT zygotes had a developed APC with a well-organised microtubule structure (Fig. 4a1-2). At this stage, multiple thread-like structures were observed within the nucleus (magenta arrowheads, Fig. 4a3-5). These structures likely represent condensed--like chromosomes (the first stage of meiotic prophase I, characterised by thread-like chromosome condensation), marking the entry into meiosis, and the ends of these chromosomes appeared to be attached to the nuclear envelope (Fig. 4a5). In contrast, *Pbnek4-ko* zygotes at 4 hpa had only small APC and microtubule structures (Fig. 4a6-7) and the condensed chromosomes seen in WT zygotes were completely absent in *Pbnek4-ko* zygotes (Fig. 4a8-10).

**Figure 4.**
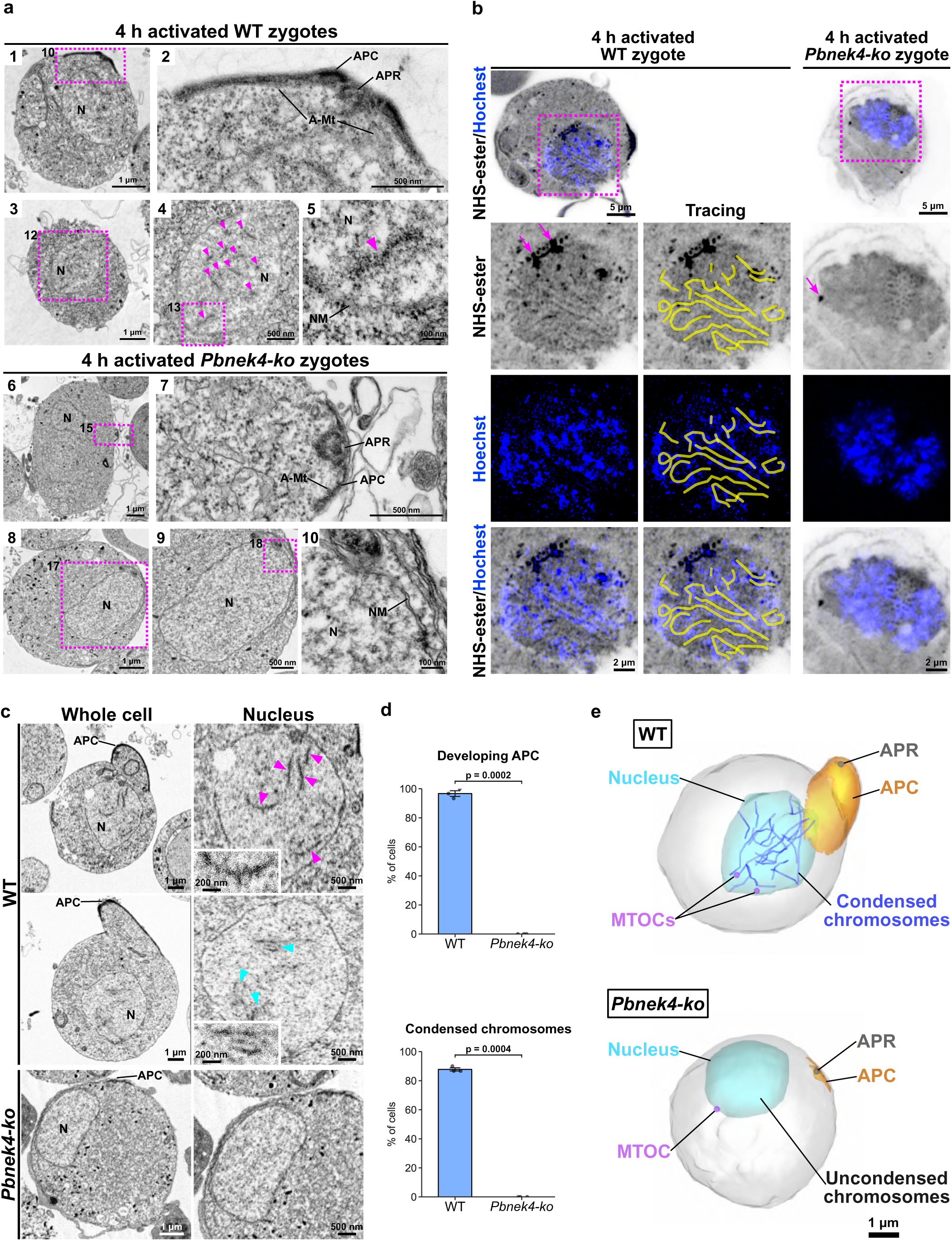
Electron microscopy shows that cytoskeletal development and condensed chromosome formation during early meiosis are blocked in *Pbnek4-ko* zygotes. **a.** TEM images of WT and *Pbnek4-ko* zygotes at 4 hpa. N, nucleus; APC, apical polar complex; APR, Apical polar ring; A-Mt, APC microtubule; NM, nuclear membrane. Magenta arrowheads show condensed chromosomes. Representative images from three independent biological experiments (n ≥ 20 cells examined per condition). **b.** U-ExM images of WT and *Pbnek4-ko* zygotes at 4 hpa, stained with NHS-ester (gray) and Hoechst (blue). A maximum intensity z-projection was generated using a subset of optical slices encompassing the nucleus (6 slices, 170-nm intervals) from the same datasets shown in Fig. 3c. Whole-cell images are shown at the top, and magnified views of the nuclear regions enclosed by magenta boxes are presented below. Magenta arrows indicate the positions of the MTOCs. Tracing images illustrate the thread-like structures within the WT nucleus traced with yellow lines based on the NHS-ester staining, demonstrating that these structural axes largely coincide with the Hoechst DNA signals. Representative images from three independent biological experiments (n ≥ 5 cells examined per condition). **c.** Slice images of WT and *Pbnek4-ko* zygotes at 4 hpa, obtained by SBF-SEM. The whole-cell view shows the development of the APC in the WT zygote. The magnified views of the nucleus show the presence of thread-like condensed chromosomes (magenta arrowheads) and the formation of potential synaptonemal complexes (cyan arrowheads), where these condensed chromosomes form pairs via a central linear structure. The insets show magnified views of a thread-like condensed chromosomes and synaptonemal complex. These developmental features were completely absent in *Pbnek4-ko* zygotes. **d.** Quantification of the percentage of cells with a developing APC and condensed chromosomes in WT and *Pbnek4-ko* zygotes at 4 hpa. The data represent the mean ± SEM from three independent SBF-SEM datasets (n = 3), in which 30 cells were analysed per dataset. with black dots representing the results from each dataset. **e.** 3D models of WT and *Pbnek4-ko* zygotes at 4 hpa, reconstructed from SBF-SEM datasets. The WT model illustrates the developed APC (orange) with APR (gray), the presence of condensed, thread-like chromosomes within the nucleus (blue; some of which are attached to the nuclear envelope), and the separating MTOCs (purple). In contrast, the *Pbnek4-ko* zygote model shows that none of these developmental events have occurred.

We also utilised U-ExM coupled with NHS-ester and Hoechst 33342 DNA staining to visualise the ultrastructure of the nucleoplasm (Fig. 4b). In WT zygotes at 4 hpa, NHS-ester staining revealed numerous distinct thread-like structures within the nucleus that colocalised with the Hoechst DNA signal, indicative of assembled chromosome axes and condensed chromosomes. In contrast, these thread-like structures were absent within the nucleus in *Pbnek4-ko* zygotes at 4 hpa, where the DNA signal remained widely diffuse.

To examine further the ultrastructural differences between WT and *Pbnek4-ko* zygotes, we used Serial Block-Face Scanning Electron Microscopy (SBF-SEM). In most WT zygotes at 4 hpa, a developed APC was detected (Fig. 4c, d) and multiple condensed, thread-like chromosomes were detected within their nuclei (Fig. 4c, magenta arrowheads and inset). In some zygotes, these chromosomes were observed pairing with linear structures between them, indicative of likely synaptonemal complex formation (Fig. 4c, cyan arrowheads and inset). In contrast, in *Pbnek4-ko* zygotes at 4 hpa there was impaired APC development, and no condensed chromosomes nor synaptonemal complexes were present within the nucleus (Fig. 4c, d). Three-dimensional models constructed from the SBF-SEM datasets illustrate the development of the APC, the formation of condensed chromosomes (with one or both ends appearing to attach to the nuclear envelope), and the separation of MTOCs in WT zygotes at 4 hpa. All these early meiotic processes are blocked in *Pbnek4-ko* zygotes (Fig. 4e). These findings indicate that the deletion of the *Pbnek4* gene severely impairs the development of the APC and its associated microtubule network in the early zygote, and also ablates the formation of condensed chromosomes (likely leptotene chromosomes)). Given that *Pbnek4-ko* parasites fail to undergo pre-meiotic DNA replication^25^, this absence of chromosome condensation is consistent with arrest prior to meiotic prophase.

### NEK4 drives meiotic entry by coordinated control of transcription, protein synthesis and phosphorylation

To elucidate the molecular processes governed by NEK4 during early meiosis in *P. berghei* zygote development, we performed an integrated transcriptomic, proteomic, phosphoproteomic, and interactomic analysis. Global transcriptomics was conducted on WT and *Pbnek4-ko* parasites at 2 hpa and quantitative proteomics/phosphoproteomics were conducted on non-activated and 2 hpa samples for both WT and *Pbnek4-ko* parasites. GFP-Trap interactomics was performed using PbNEK4-GFP parasites at 2 hpa.

Quantitative real-time PCR (qRT-PCR) showed that the expression of meiosis-related genes (e.g., *dmc1*, *hop1*, *mnd1*, and *spo11*) was significantly downregulated in *Pbnek4-ko* zygotes at 2 hpa compared to WT zygotes at 2 hpa (Extended Data Fig. 6a). Transcriptomic profiling via RNA-sequencing between WT and *Pbnek4-ko* parasites at 2 hpa revealed major transcriptional changes in *Pbnek4-ko* parasites. At 2 hpa, 1,554 genes were significantly differentially expressed (adjusted p-value ≤ 0.05, fold change ≥ 2) relative to WT parasites, comprising 717 downregulated and 837 upregulated genes (Fig. 5a; Supplementary Table 1). Gene Ontology enrichment of downregulated transcripts highlighted key biological processes, including cell motility, cell differentiation, meiotic nuclear division, and cytoskeletal organisation – hallmarks of zygote maturation (Fig. 5b; Supplementary Table 2). Notably, transcripts encoding the meiotic recombinase *dmc1* ^39^ and the putative synaptonemal complex component *hop1* (PBANKA_1407900)^40^ were strongly reduced in the *Pbnek4-ko* line.

**Figure 5.**
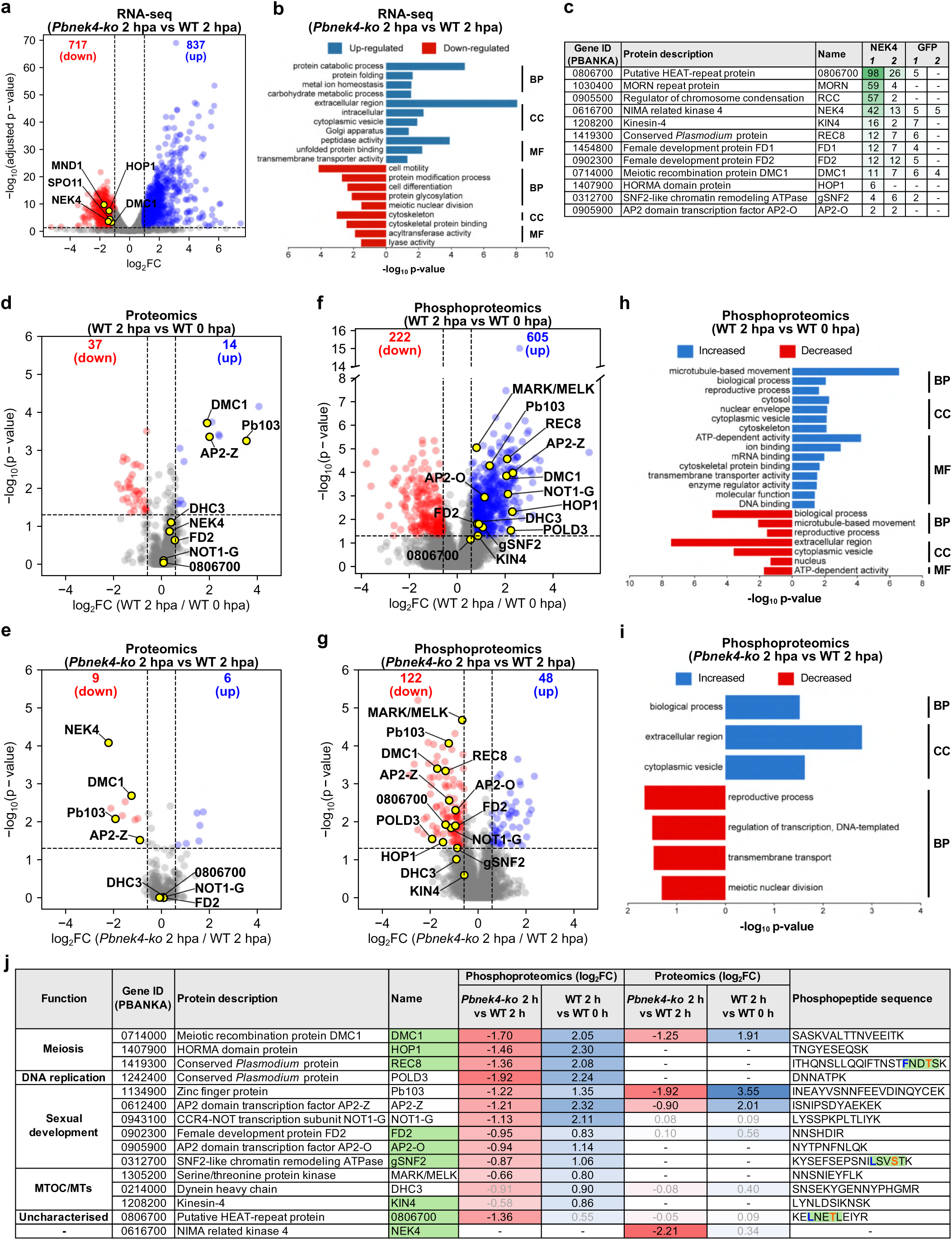
Global transcriptomic, proteomics and phosphoproteomic analysis reveals NEK4-dependent transcriptional and phosphorylation events during *Plasmodium* zygote activation. **a.** Volcano plot showing differentially expressed transcripts in *Pbnek4-ko* parasites compared to WT controls at 2 hpa (RNA-seq). The log_2_ fold change (FC) is plotted against -log_10_(adjusted p-value). Red and blue dots indicate significantly upregulated and downregulated genes, respectively. *NEK4* and key meiotic genes that are significantly downregulated in *Pbnek4-ko* at 2 hpa are highlighted with yellow dots and gene names. Dotted lines indicate the threshold for statistical significance (adjusted p-value ≤ 0.05 and fold change ≥ 1.5). The underlying data are provided in Supplementary Table 1. **b.** Gene Ontology (GO) analysis of differentially expressed transcripts by RNA-seq in *Pbnek4-ko* versus WT zygotes at 2 hpa. Genes in each term are listed in Supplementary Table 2. The analysis was performed using GO slim terms in PlasmoDB. BP = Biological Process; CC = Cellular component; MF = Molecular Function. **c**. Table displaying selected proteins, corresponding gene ID and representation by the number of peptides from the proteins identified in a GFP-trap pulldown assay with precipitates of either PbNEK4-GFP or WT-GFP zygotes at 2 hpa. **d-g.** Volcano plots displaying changes in total protein and phosphopeptide abundance. The log_2_ fold change (derived from the average of three biological replicates each) is plotted against the -log_10_ p-value. The dotted lines indicate the threshold for statistical significance (p-value ≤ 0.05 and fold change ≥ 1.5). Significantly upregulated and downregulated proteins/phosphopeptides are shown as red and blue dots, respectively. Proteins/phosphopeptides with non-significant changes are shown as grey dots. **d.** Comparison of protein abundance between WT gametocytes at 0 hpa and WT zygotes at 2 hpa. **e.** Comparison of protein abundance between *Pbnek4-ko* and WT zygotes at 2 hpa. **f.** Comparison of phosphopeptide abundance between WT gametocytes at 0 hpa and WT zygotes at 2 hpa. **g.** Comparison of phosphopeptide abundance between *Pbnek4-ko* and WT zygotes at 2 hpa. Key phosphopeptides from proteins of interest that are significantly upregulated in WT at 2 hpa (f) and downregulated in the *Pbnek4-ko* mutant at 2 hpa (g) are highlighted with yellow dots and protein names. **h-i.** GO analysis of differentially abundant phosphopeptides in WT zygotes at 2 hpa versus 0 hpa (h) and *Pbnek4-ko* versus WT zygotes at 2 hpa (i). Genes in each term are listed in Supplementary Table 8 and 10. The analysis was performed using GO slim terms in PlasmoDB. **j.** Summary table displaying selected proteins with significant changes in phosphopeptide abundance from the phosphoproteomics screen. Protein names highlighted with a green background indicate that these proteins were also independently identified in the PbNEK4-GFP pulldown. The log_2_ fold change (log_2_FC) values of identified phosphopeptides are represented by a colour gradient from red (decrease) to blue (increase). Significant log_2_FC values (p-value ≤ 0.05) are shown in black text, while non-significant values are in grey. In the phosphopeptide sequences, the regions corresponding to the canonical NEK kinase motif ([LMFW]-X-X-S/T-[no P])^55^ are highlighted with a green background. Within these motifs, the hydrophobic residues at the -3 position are indicated in blue font, and the target phosphorylated serine/threonine residues are indicated in orange font.

Next, we performed GFP pulldown assays using PbNEK4-GFP and WT-GFP zygotes at 2 hpa. This analysis identified interactions of PbNEK4-GFP with putative MTOC-associate proteins, such as a putative HEAT-repeat protein, which possesses predicted HEAT-like repeats revealed by AlphaFold3^41^ and Foldseek^42^ structural prediction (PBANKA_0806700) and a MORN repeat protein (PBANKA_1030400), as well as a regulator of chromosome condensation (RCC; PBANKA_0905500). Furthermore, interactions with meiotic factors, including REC8 (PBANKA_1419300)^43^, DMC1, and HOP1, were also indicated (Fig. 5c; Supplementary Table 13). These interactions may suggest that NEK4 acts as a signalling hub, and either directly phosphorylates or stabilises other components within an integrated meiotic complex.

To investigate the full landscape of NEK4-associated protein abundance and phosphorylation changes during early zygote development, we performed quantitative proteomics and phosphoproteomics analyses using non-activated (0 hpa) WT and *Pbnek4-ko* gametocytes, and WT and *Pbnek4-ko* zygotes at 2 hpa. This design enabled us to separate activation-associated changes (WT 2 hpa vs WT 0 hpa) from NEK4-dependent effects during early zygote development (*Pbnek4-ko* 2 hpa vs WT 2 hpa). Initial analysis of phosphopeptide-enriched and flow-through fractions from *Pbnek4-ko* and WT gametocytes at 0 hpa revealed minimal differences in the abundance of identified proteins and phosphopeptides (Extended Data Fig. 6b and d; Supplementary Table 5 and 12). These findings indicate that *Pbnek4* deletion does not substantially reprogramme the proteome or phosphoproteome in mature gametocytes, and instead suggest that NEK4-dependent changes are initiated upon activation/fertilisation.

In contrast, significant differences in protein and phosphopeptide abundance were observed when comparing gametocytes at 0 hpa with zygotes at 2 hpa, and *Pbnek4-ko* with WT zygotes at 2 hpa (Fig. 5d-g; Extended Data Fig. 6c and f; Supplementary Table 6 and 11). Proteomic analysis of the flow-through fractions obtained during phosphopeptide enrichment revealed marked increases in zygote- and meiosis-associated proteins following fertilisation in WT parasites. These included the zinc-finger protein Pb103^44^, the transcription factor AP2-Z^45^, and the meiotic recombinase DMC1^39^ (Fig. 5d; Supplementary Table 3). In contrast, when *Pbnek4-ko* parasites were compared to WT controls at 2 hpa, many of these same proteins were significantly less abundant (Fig. 5e; Supplementary Table 4), suggesting that NEK4 activity influences their synthesis, stability, or translation during zygote development.

Phosphoproteomic profiling revealed a similar association with the presence of NEK4 for protein phosphorylation linked to meiotic initiation. In WT parasites, fertilisation triggered widespread phosphorylation of proteins linked to meiosis, transcription, translation, and cytoskeletal organisation by 2 hpa. These proteins included DMC1^39^, HOP1^40^, REC8^43^, DNA polymerase delta subunit 3 (POLD3), AP2-Z^45, 46^, AP2-O^47^, Pb103^44^, FD2^48^, gSNF2^49^, NOT1-G^50^, a putative HEAT-repeat protein (PBANKA_0806700), a serine/threonine protein kinase putatively related to MAP/microtubule affinity-regulating kinase 1^51^ and maternal embryonic leucine zipper kinase 1^52^ (MARK/MELK), zygote-ookinete specific dynein heavy chain (DHC3)^53^, and kinesin-4 (KIN4)^54^ (Fig. 5f; Supplementary Table 5). In *Pbnek4-ko* parasites phosphorylation at many sites in these proteins was significantly lower relative to the WT control (Fig. 5g; Supplementary Table 9), Notably, some of these proteins also showed increased overall abundance in the flow-through fraction (e.g. DMC1, Pb103, and AP2-Z), indicating that the observed phosphorylation changes may partly reflect higher protein levels rather than phosphorylation alone. However, the consistent reduction of phosphorylation in *Pbnek4-ko* parasites suggests that NEK4 contribute to phosphoregulation in addition to promoting protein accumulation. Gene Ontology (GO) analysis further confirmed that zygote activation triggers increased phosphorylation of proteins involved in microtubule-based movement, the cytoskeleton, and reproductive processes (Fig. 5h; Supplementary Table 8). In contrast, *Pbnek4-ko* zygotes at 2 hpa showed reduced phosphorylation of proteins associated with reproductive processes and meiotic nuclear division (Fig. 5i; Supplementary Table 10). Representative proteins identified by our proteomics and phosphoproteomic analyses, along with their abundance and putative functions, are summarised in Figure 5j. These data support the hypothesis that NEK4 may contributes to a phosphorylation network underpinning zygote differentiation and meiotic progression.

Mammalian NEK kinases favour a hydrophobic residue (Leu, Met, Phe, or Trp) at the -3 position relative to the target serine or threonine, and disfavour a proline residue at the +1 position, establishing a consensus motif of [LMFW]-X-X-[S/T]-[no P]^55, 56^. We observed this consensus motif in multiple high-confidence *Plasmodium* targets ([F]-N-D-[T]-[S] in REC8, [L]-S-V-[S]-[T] in gSNF2, and [L]-N-E-[T]-[L] in PBANKA_0806700). Importantly, all of these motif-containing proteins were also independently identified as physical interaction partners in the NEK4 GFP-Trap interactome (Fig. 5c and j).

Collectively, these integrated transcriptomic, interactomic, proteomic, and phosphoproteomic datasets suggest that while NEK4 may exert some influence on transcript and protein accumulation, its most profound and immediate impact at this early development stage (2 hpa) is on the phophoproteome. Together with the interactomic evidence, these results identify NEK4 as a key regulator of early post-zygotic meiosis in *P. berghei*, orchestrating a rapid rewiring of phosphosignalling networks to drive meiotic entry and zygote maturation.

## Discussion

We identify *Plasmodium*NEK4 as a key kinase that links meiotic initiation and zygote morphogenesis in *P. berghei* (Fig. 6). NEK4 localises to both the MTOC and the APC during the earliest stages of zygote development (post-zygotic meiosis), and orchestrates MTOC duplication, perinuclear and APC microtubule formation, and migration of the nucleus. This directed movement resembles the “horsetail” nuclear oscillations during the meiotic prophase in other eukaryotes, where cytoskeleton-driven repositioning of the nucleus promotes homologous chromosome pairing and recombination^32, 37^. In *S. pombe*, this nuclear movement occurs during prophase, where linear elements (LinEs) mediate chromosome pairing and crossover. Similarly, in *Plasmodium*, from ∼4 hpa onwards —when DNA replication is complete, nuclear movement is evident, and morphological transformation into the ookinete begins —the formation of condensed chromosomes reminiscent of leptotene chromosomes and structures resembling synaptonemal complexes were observed. Although fully condensed metaphase chromosomes seen in canonical meiosis are still elusive in *Plasmodium*, this initial condensation step is clearly a prerequisite for proper meiotic progression and this suggests that entry into meiotic prophase initiates at this stage in *Plasmodium* (Fig. 6). In the absence of NEK4, the MTOC fails to duplicate, chromatin remains diffuse, and nuclear translocation is lost, leading to arrest of cell development before meiotic division and transformation to ookinete. These data suggest that NEK4 is a putative central regulator linking nuclear reorganisation and cytoskeletal polarity at the onset of meiosis. Previous studies suggest that while *Plasmodium* retains some canonical meiotic processes well-characterised in organisms like *S. pombe*, its mechanisms and molecular players are highly divergent, as evidenced by the evolutionary conservation analysis of meiotic components in *Plasmodium* and by the progression through meiosis I and II without nuclear division^8^ (Fig. 6). Given this divergence, it is particularly intriguing that early meiotic nuclear movement observed in model eukaryotes is conserved in *Plasmodium*, suggesting that it may be important for homologous chromosome pairing and recombination in this parasite as well.

**Figure 6.**
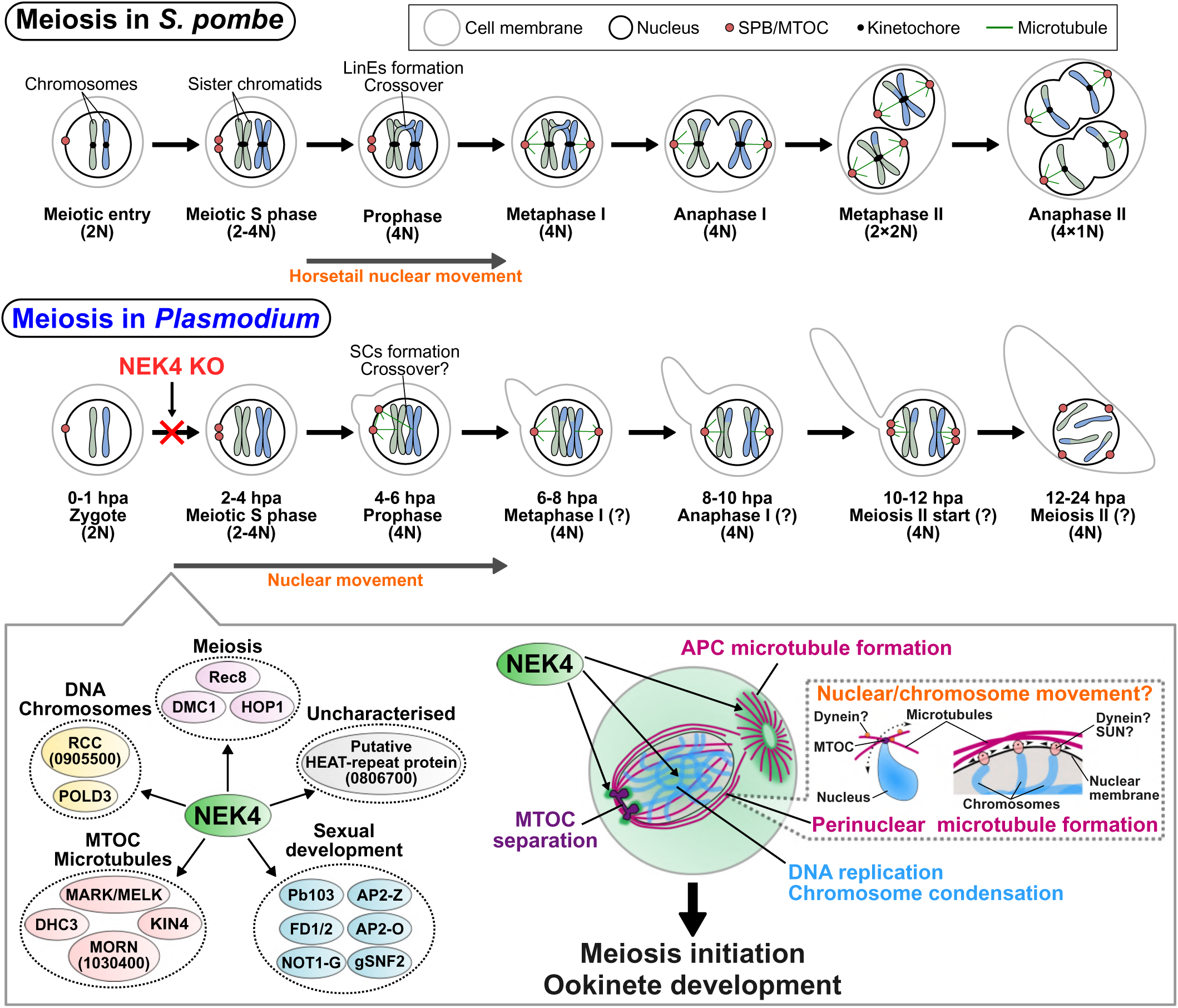
*Plasmodium* NEK4 acts as a key regulator driving the early events of meiosis. A model illustrating the central function of NEK4 in coordinating key developmental processes during the zygote-to-ookinete development. The top panel shows the canonical meiotic stages in the model organism *Schizosaccharomyces pombe* (*S. pombe*) for comparison. The middle panel depicts the predicted steps and timing of meiosis in *Plasmodium* from zygote formation, meiosis I to meiosis II, based on previous observations. In *Pbnek4-ko* mutants, DNA replication is impaired, blocking progression to meiotic prophase. In *S. pombe*, active nuclear movement, known as horsetail nuclear movement, occurs during prophase, followed by chromosome pairing and crossover mediated by linear elements (LinEs). In contrast, our observations in *Plasmodium* suggest that between 2 and 4 hpa, active nuclear movement similar to the horsetail movement occurs concurrently with the formation of structures resembling synaptonemal complexes between condensed chromosomes. The bottom panel proposes that NEK4 regulates four crucial modules: (1) MTOC/Microtubule dynamics (e.g. MARK/MELK, MORN), (2) Meiosis-specific proteins (e.g. REC8, HOP1), (3) Sexual development factors (e.g., FD2, AP2-O), (4) DNA/chromosome functions (e.g. RCC, POLD3), (5) Uncharacterised protein (putative HEAT-repeat protein; PBANKA_0806700). Through these pathways, NEK4 orchestrates MTOC separation, microtubule formation at MTOC and APC, DNA replication, and chromosome condensation, which are all essential for successful meiosis and subsequent ookinete development. Based on the function of perinuclear microtubules in other eukaryotes during meiosis, the perinuclear microtubules observed in *Plasmodium* zygotes are likely associated with nuclear movement or intranuclear chromosome movement. We predict that effectors such as cytoplasmic dynein and SUN protein mediate these functions.

The dual location of NEK4 and the developmental arrest that results from knocking out its gene, place it at the intersection of nuclear and cytoskeletal organisation. Its behaviour parallels that of NEKs in other systems, which regulate centrosome duplication, spindle formation, and checkpoint transitions^22, 23^. In *S. pombe*, the NEK-like kinase Fin1 controls spindle formation, specifically the maturation and inheritance of the SPB^57, 58^, while NEK2 and NEK9 act at metazoan centrosomes to coordinate spindle assembly^23^. In *Plasmodium*, which lacks canonical mitotic and meiotic regulators such as the phosphatases CDC25, CDC14, and Polo-like kinases^18, 19^, NEK4 appears to have acquired functions to integrate both signalling and structural control within a simplified regulatory network.

Proteins identified in the PbNEK4-GFP pulldown assays, such as the putative HEAT-repeat protein (PBANKA_0806700), MORN repeat protein (PBANKA_1030400), and Kinesin-4 (PBANKA_1208200), may interact with NEK4 at the MTOC to facilitate MTOC development or segregation. Although the specific function of PBANKA_0806700 in *Plasmodium* remains uncharacterised, HEAT-repeat proteins are universally known to form alpha-helical solenoids that act as versatile scaffolds for protein-protein interactions^59^. Given its massive size (4,658 aa) and multiple predicted HEAT repeats, we hypothesise that this protein serves as a central structural scaffold at the MTOC, organising the macromolecular complexes required for NEK4-dependent cytoskeletal remodelling during early zygote development. Additionally, the identified regulator of chromosome condensation (RCC; PBANKA_0905500) contains an RCC1 domain. While RCC1 domains classically mediate interactions with chromatin or DNA^60^, their functions are not restricted to chromosomal regulation. Interestingly, human NEK8 and NEK9 possess an RCC1 domain, and this domain is implicated in the recruitment to the centrosome^61^. Furthermore, a recent study in *Toxoplasma gondii* has demonstrated that RCC-domain-containing proteins are involved in the apical complex assembly^62^. This raises the possibility that the *Plasmodium* RCC identified here is similarly recruited to the MTOC or the APC, where it may interact with NEK4 to coordinate cytoskeletal and apical organisation.

Transcriptomic, proteomic and phosphoproteomic profiling revealed that fertilisation triggers extensive reprogramming of transcription, protein abundance, and protein phosphorylation during post-zygotic meiosis. While the apparent downregulation of many factors between 0 and 2 hpa is heavily confounded by the natural depletion of male gametocytes following activation, our targeted comparison between WT and *Pbnek4-ko* at 2 hpa successfully uncoupled these effects to reveal the specific NEK4-dependent networks. In WT parasites, proteins likely essential for meiosis and zygote differentiation including protein sets relating to meiosis (DMC1, HOP1 and REC8), DNA replication (POLD3), sexual development (Pb103, AP2-Z, NOT1-G, FD2, AP2-O, gSNF2), and MTOC/microtubules (MARK/MELK, DHC3, KIN4) are more phosphorylated in the zygote (Fig. 6). In *Pbnek4-ko* parasites, these proteins were hypophosphorylated, suggesting that NEK4 contributes to phosphorylation events required for onset of meiosis and zygote-ookinete development. Also, while the primary effect of *Pbnek4* deletion is a sweeping reduction in phosphorylation, the concomitant reduction in protein abundance for a subset of specific factors (e.g. DMC1, Pb103, and AP2-Z) suggests that NEK4-dependent phosphosignalling may secondarily influence the stability or synthesis of certain key meiotic regulators.

While distinguishing direct kinase substrates from downstream cascading effects is inherently challenging in knockout studies, our integrated multi-omics datasets provide a powerful filter. Analysis of the specific hypophosphorylated sites identified in our phosphoproteomic dataset reveals that several proteins - such as REC8, gSNF2, and the putative HEAT-repeat protein PBANKA_0806700 - possess sequence motifs that perfectly align with canonical NEK kinase preferences. Mammalian NEK kinases favour a hydrophobic residue (Leu, Met, Phe, or Trp) at the -3 position relative to the target serine or threonine, and disfavour a proline residue at the +1 position, establishing a consensus motif of [LMFW]-X-X-[S/T]-[no P]^55, 56^.

We observed this structural signature in multiple high-confidence *Plasmodium* targets ([F]-N-D-[T]-[S] in REC8, [L]-S-V-[S]-[T] in gSNF2, and [L]-N-E-[T]-[L] in PBANKA_0806700). Strikingly, all of these motif-containing proteins were also independently identified as physical interaction partners in our NEK4 GFP-Trap interactome. The convergence of canonical motif-dependent phosphorylation and physical interaction strongly supports their identity as likely substrates of NEK4. Although the limited number of these validated sites currently precludes the mathematical definition of a robust, *de novo* PbNEK4 consensus motif, these findings provide a critical molecular starting point for understanding how NEK4 directly interfaces with the early meiotic and developmental machinery.

Such coordinated phosphoregulation is consistent with broader concepts being developed for eukaryotic meiosis. Recent phosphoproteomic studies in yeast have shown that meiotic progression is driven by both changes in protein abundance and dynamic, stage-specific waves of protein phosphorylation that rewire existing signalling networks^63^. These phosphorylation cascades, directed by cyclin-dependent and Polo-like kinases, define temporal transitions from recombination to segregation^63^. Large-scale quantitative phosphoproteomics studies in *S. pombe* revealed that phosphorylation of thousands of sites changes during meiotic development, for example affecting proteins involved in microtubule organisation, recombination and chromosome cohesion^64^. Our findings are consistent with this concept of a meiotic “phosphorewiring”: the absence of NEK4 collapses a broad phosphorylation programme encompassing transcriptional, translational and cytoskeletal effectors, and other kinases and phosphatases, and implying that *Plasmodium* has an analogous, temporally-ordered kinase programme despite its highly reduced kinase signalling repertoire.

Although our study focuses on NEK4-dependent phosphorylation and protein accumulation during early zygote development, the quantitative phosphoproteomics also identify multiple protein kinases and phosphatases with substantial phosphorylation changes across activation and/or in *Pbnek4-ko* parasites, and several kinase/phosphatase proteins are additionally detected in the NEK4-GFP interactome (Supplementary Tables 13 and 14). In our curated view of the dataset, phosphorylation of a PP2B/calcineurin catalytic subunit (CNA; PBANKA_1227400), several PPM-family serine/threonine phosphatases (including PPM5/6/8/11), and an NIF1-like phosphatase are dynamically modified during the 0–2 hpa transition, with at least one PPM-family phosphatase (PPM5; PBANKA_1427200) showing reduced phosphorylation in *Pbnek4-ko* relative to WT at 2 hpa. These observations suggest that NEK4-associated “rewiring” is likely to involve not only kinase-driven phosphorylation cascades but also changes in phosphatase activity, recruitment, or substrate access. The presence of several phosphatases among the NEK4-dependent phosphosites and interactors suggests that NEK4 participates in a broader signalling network balancing kinase and phosphatase activities during early zygote development. Similar kinase–phosphatase regulatory modules are known to control meiotic transitions in other eukaryotes, including the regulation of chromosome segregation and meiotic progression by PP2A-family phosphatases^65, 66^. Notably, conserved phosphatase control is also implicated in *Plasmodium* atypical division programmes^18^, supporting the possibility that phosphatases contribute significantly to parasite signalling during sexual development.

Additionally, our datasets highlight the early zygote transcription factor AP2-Z as a NEK4-associated node, with altered abundance and reduced phosphorylation in *Pbnek4-ko* parasites at 2 hpa. Analysis of the transcriptional changes in *Pbnek4-ko* zygotes at 2 hpa showed that ∼41% of annotated AP2-Z targets and ∼41% of AP2-O targets, respectively are significantly altered in the *Pbnek4-ko* line (Supplementary Tables 15 and 16), consistent with a broad disruption of the stage-specific transcriptional outputs normally associated with these AP2 regulators during zygote/ookinete development. In contrast, when focusing specifically on canonical meiotic genes, DMC1 was the only factor among those highlighted in our study and is a reported AP2-Z/O target^45, 67^. This suggests that NEK4-dependent control of meiotic initiation is not explained just by direct AP2-Z/O regulation of meiotic gene transcription but may also involve post-transcriptional regulation and phosphosignalling-dependent changes in protein accumulation.

Little is known about phosphorylation-dependent activation of AP2-Z or AP2-O. The transcription of AP2-Z is regulated by upstream gametocyte factors (AP2-G and AP2-FG) binding to the AP2-Z promoter^68^. The AP2-Z/O phosphosite(s) in the phosphoproteomic datasets lie outside the AP2 DNA-binding domain, but this does not preclude a regulatory role; the phosphorylation may modulate interactions with co-factors or chromatin-associated complexes, affect stability, or determine subcellular localisation. These features may be particularly important in the tightly staged zygote-to-ookinete transcriptional transition. Together, these findings support a model in which NEK4 contributes to early developmental “rewiring” that intersects with AP2-led transcriptional programmes. Defects in meiotic progression likely reflect a combination of contributions from transcription, translation and phosphoregulation.

It is important to note a biological limitation to our analyses: the 0 hpa parasite population is comprised of a mixture of both male and female gametocytes. Following activation, male gametes exflagellate and their biomass is largely depleted from the developing zygote culture. Consequently, a substantial proportion of the proteins and phosphosites that are diminished at 2 hpa (including known male-specific factors such as MDV1) likely reflect this shift rather than active developmental downregulation within the female/zygote lineage. Additional sampling earlier than at 2 hpa is likely to further enrich for primary NEK4-dependent phosphorylation and interactions, and will be important for resolving the temporal order of the kinase/phosphatase activation and inhibition underlying the rewiring.

Our structural observations suggest that the APC and the microtubules extending from the nuclear MTOC are formed in a NEK4-dependent manner. Furthermore, we observed a link between the microtubules extending from the MTOC and APC, suggesting that this connection is associated with nuclear movement or positioning. Comparable architectures are present in other apicomplexans: in *Toxoplasma gondii*, the APC and conoid form an anchoring hub that coordinates cortical microtubules and organelle positioning^69, 70^. *Eimeria tenella* and *Cryptosporidium* spp. also have similar apical scaffolds^71, 72^. Genetic modification of key regulatory kinases has resulted in comparable defects in other apicomplexans-specifically, the failure of MTOC/centrosome duplication and the uncoupling of nuclear replication from cytoskeletal assembly—supporting a conserved requirement for precise coordination between nuclear and cortical assembly. In *T. gondii*, conditional mutants of NEK1 and the MAP kinase, MAPK2 block centrosome duplication and segregation, causing the nucleus to replicate without successful daughter cell budding, effectively decoupling nuclear and cytoskeletal programmes^73, 74^. *Plasmodium* NEK1 controls spindle and kinetochore organisation during male gametogenesis^28^, while loss of the phosphatase PPKL disrupts apical complex integrity and ookinete morphogenesis^18, 75^. A putative MARK/MELK-related serine/threonine protein kinase (PBANKA_1305200), identified in our phosphoproteomics analysis, may be directly or indirectly activated by NEK4 to stimulate microtubule-associated proteins, thereby promoting or stabilising microtubule formation at the APC and MTOC in the zygote. Together, these studies highlight a recurrent feature of Apicomplexa: kinases and phosphatases govern centrosome and apical organisation, and their perturbation uncouples nuclear and cortical events, resulting in developmental arrest. The function of NEK4 fits within this broader mechanistic framework, acting as a linchpin to preserve synchrony between meiotic and morphogenetic processes.

The perinuclear microtubules observed in early zygotes may be involved in the nuclear movement described above and intranuclear chromosome movement (Fig. 6). In *S. pombe*, horsetail nuclear movement is driven by SPB-derived microtubules interacting with the cell cortex via dynein^34, 35^. We speculate that a similar mechanism generates nuclear movement in *Plasmodium*, through the interaction of perinuclear microtubules with dynein (e.g, DHC3, which was identified in our phosphoproteomics). Indeed, our preliminary observation that PbDHC3 is largely located at the nuclear periphery in early zygotes strongly supports this hypothesis (Extended Data Fig. 7 and Supplementary Video 8 and 9). While determining the precise biophysical mechanics and the biological necessity of this movement for meiotic completion will require future functional studies utilising dynein inhibitors and high-resolution tracking of microtubule dynamics, the recruitment of DHC3 provides a compelling molecular candidate for driving this nuclear motility. Furthermore, in eukaryotes including mice, nematodes, and yeast, it is well established that meiotic chromosomes interact with dynein on perinuclear microtubules via the SUN (Sad-1 and UNC-84)/KASH (Klarsicht, ANC-1, and SYNE Homology) complex on the nuclear envelope, facilitating chromosome movement and pairing^76^. Given that SUN proteins^77^ and DHC3 localise to the nuclear periphery during the zygote-to-ookinete development in *Plasmodium*, it is plausible that SUN proteins mediate the interaction between chromosomes and dynein on perinuclear microtubules, thereby driving intranuclear chromosome movement to promote homologous chromosome pairing.

In an evolutionary context, NEK4 exemplifies how *Plasmodium* has streamlined but not abandoned a central feature of meiotic control, to ensure synchrony between nuclear and cytoskeletal events. The parasite has a compact kinome, and a limited number of four NEKs have diversified to perform additional discrete yet complementary roles: for example, NEK1 regulates spindle formation during male gametogenesis^28^, while NEK4 governs meiotic initiation and zygote differentiation. This division of labour underscores the adaptive flexibility of NEKs in the absence of canonical Polo-kinases and other regulators. The NEK4 regulatory module may be a *Plasmodium*-specific version of an ancient, conserved solution to a universal cellular problem – to use specific waves of phosphorylation to coordinate chromosomal replication with structural reorganisation of the cell during meiosis.

In conclusion, NEK4 functions as a key regulator that co-ordinates meiotic entry in *Plasmodium*, coupling microtubule organisation, MTOC duplication, nuclear migration, DNA replication, and chromosome condensation with the establishment of apical polarity (Fig. 6). By integrating translational and phosphoregulatory control, NEK4 contributes to the coordination of nuclear and cytoskeletal events, which is essential to drive zygote morphogenesis and meiosis. These findings clarify the molecular requirements of meiosis in Apicomplexa, highlighting a conserved principle – kinase-mediated coordination of nuclear and cellular processes – that extends across the eukaryotic kingdom. Given that NEK4 is essential for the development of transmission-stage parasites and the absence of close orthologs from the human host, this kinase represents a promising transmission-blocking therapeutic target. NEK4 provides a window to investigate how the minimalist signalling networks of divergent parasites sustain their complex developmental programmes.

## Methods

### Ethics statement

All animal experiments were conducted in the United Kingdom in accordance with the Animals (Scientific Procedures) Act 1986 and approved by the Home Office under Project Licence numbers PDD2D5182 and PP3589958. Protocols were reviewed and approved by the institutional Animal Welfare and Ethical Review Body (AWERB) prior to implementation. Procedures were designed to minimise animal suffering and the number of animals used, in line with the 3Rs (Replacement, Reduction, Refinement) principles. Experiments were performed on female CD1 outbred mice aged 6 to 8 weeks, housed under standard conditions with environmental enrichment and monitored daily for health and welfare.

### Generation of transgenic parasites

For C-terminal GFP-tagging of NEK4 (PBANKA_0616700) by single crossover homologous recombination, a 930 bp region of *nek4* downstream of the ATG start codon was amplified using primers 5’-CCCCGGTACCGATAGCGATAAGAGAGTAAGATTGTGTG-3’ and 5’-CCCCGGGCCCAACATCAACAATATCCAATAATAATG-3’ (Supplementary Table 17). Genotype analysis was performed using integration PCR as illustrated in Extended Data Fig. 1d. Primer 1 (intNEK4: 5’-CCAACCATATATTACACAGAGGTTAG-3’) and primer 2 (ol492)^18^ were used to confirm correct integration of the *gfp* sequence at the target locus. *P. berghei* ANKA line 2.34 parasites were then transfected by electroporation^78^; confirmation of the correct protein size was performed by Western blot analysis^28^. Production of the *Pbnek4-ko* parasite line has been described previously^19, 25^.

### Live cell imaging

To analyse PbNEK4-GFP expression during the zygote to ookinete transition, gametocytes or zygotes activated in ookinete culture medium were purified by Nycodenz gradient^19^. Zygote and ookinete development were maintained in continuous culture for up to 24 hours post-fertilisation. At specific time points, cell aliquots were taken and immediately mixed with a Cy3-conjugated mouse monoclonal antibody 13.1 (which targets the P28 surface protein)^79^ and Hoechst 33342 prior to live-cell imaging. This approach ensured that the images represent distinct, representative live cells at each time point, by avoiding phototoxicity or non-physiological effects associated with prolonged exposure to the antibody and the DNA dye. Zygote and ookinete development were tracked for up to 24 hours post-fertilisation to assess the dynamics of PbNEK4-GFP location during meiotic progression and morphogenesis. Images were captured on a Zeiss Axio Imager M2 microscope equipped with a 63× oil immersion objective and an AxioCam ICc1 digital camera (Carl Zeiss). Fluorescence signals from GFP and Hoechst were prone to bleaching, but imaging conditions were optimised to ensure sufficient resolution to follow protein dynamics across key developmental transitions. Axiovision software (Rel. 4.8) was used to enhance contrast and minimise background, ensuring threshold settings remained consistent across samples. Representative images from at least three independent biological experiments are presented, with protein location assessed in 30–50 individual cells per time point. For the quantification of PbNEK4-GFP localisation, data were derived from a total of 30–50 individual cells per time point, collected across at least three independent biological experiments. To account for variations in the microscopic focal plane (Z-axis) which can obscure simultaneous observation of different cellular regions, cells were independently scored for the presence of a strong/distinct focus at the nucleus and at the cell periphery. The percentage of cells positive for distinct focal signals at each location was calculated and plotted.

### Transmission electron microscopy

For ultrastructural analysis of NEK4-deficient and wild-type parasites, zygotes were harvested at 2 and 4 hpa and fixed in 4% glutaraldehyde in 0.1 M phosphate buffer. Samples were processed for transmission electron microscopy following the protocol described in the previous study^80^. Briefly, fixed cells were post-fixed in osmium tetroxide, stained en bloc with uranyl acetate, dehydrated through a graded ethanol series, and embedded in Spurr’s epoxy resin. Ultrathin sections (approximately 70 nm) were cut using a diamond knife, mounted on copper grids, and stained with uranyl acetate followed by lead citrate to enhance contrast. Sections were examined using a Tecnai G2 12 BioTwin (FEI UK, UK) or a JEOL 1200EX transmission electron microscope (JEOL, Japan) operated at 100 kV. Representative images were captured digitally for analysis.

### Serial block face scanning electron microscopy (SBF-SEM) of zygotes at 4 hpa

Samples were processed for SBF-SEM following the protocol described previously^81^. Fractions enriched in WT and *Pbnek4-ko* zygotes at 4 hpa were fixed at room temperature in 4% glutaraldehyde in 0.1□M phosphate buffer. Samples were then spun and washed three times in 0.1□M phosphate buffer, and post-fixed in 1% osmium tetroxide in 1.5% potassium ferrocyanide in 0.1□M phosphate buffer (for 45□min at room temperature, and in the dark). After the first osmium step, samples were washed three times in 0.1□M phosphate buffer, incubated in 1% tannic acid in 0.1□M phosphate buffer for 30□min at room temperature, and then subjected to a second osmium step (2% OsO_4_ in ddH_2_O, for 30□min, at room temperature, and in the dark). Samples were then incubated in 2% uranyl acetate in ddH_2_O for 2□h, dehydrated in acetone (a progressive series of acetone concentrations from 20%, 40%, 90% to 100% with three changes in molecular-sieved ultradry acetone over 4□h) and embedded in TAAB 812 Hard resin (TAAB, catalogue number T030). The tips of resin blocks containing samples were trimmed and mounted onto aluminium pins using conductive epoxy glue and silver dag, and then sputter coated with a layer (10–13□nm) of gold, in an Agar Auto Sputter Coater (Agar Scientific). Before SBF-SEM imaging, ultrathin sections (70□nm) of the block face were examined in a Jeol JEM 1400 Flash transmission electron microscope (JEOL), to verify sample quality. Samples were then imaged in a Merlin VP compact high resolution scanning electron microscope (Zeiss) equipped with a 3View stage (Gatan-Ametek), and an OnPoint back-scattered electron detector (Gatan-Ametek), in variable pressure. The following imaging conditions were used: 2□kV, 20□µm aperture, 30 pascal variable pressure, 2□nm pixel size, 5□μs pixel time, 100□nm section thickness, 100% FCC.

### Ultrastructure expansion microscopy

Ultrastructure expansion microscopy (U-ExM) was performed on PbNEK4-GFP non-activated gametocytes or zygotes at 2 or 4 hpa, following previously published protocols with slight modifications^82, 83^. Parasites were fixed in 4% formaldehyde in MTSB buffer (10 mM MES, 150 mM NaCl, 5 mM EGTA, 5 mM MgCl_2_, 5 mM glucose, pH7.0) and adhered to 10 mm poly-D-lysine–coated round coverslips for 15 min. Coverslips were incubated overnight at 4°C in 1.4% formaldehyde (FA)/2% acrylamide (AA). Gelation was performed in ammonium persulphate/TEMED (10% each)/monomer solution (23% sodium acrylate; 10% AA; 0.1% BIS-AA in PBS) on ice for 5 min and at 37°C for 30 min. Gels were denatured for 15 min at 37°C and for 90 min at 95°C in denaturation buffer (200 mM SDS, 200 mM NaCl, 50 mM Tris, pH 9.0, in water). After denaturation, gels were incubated in distilled water overnight for complete expansion. The next day, circular gel pieces with a diameter of ∼13 mm were excised, and the gels were washed in PBS three times for 15 min to remove excess water. The gels were then incubated in blocking buffer (3% BSA in PBS) at room temperature for 30 min, incubated with rabbit polyclonal anti-GFP antibody (1:250 dilution; A11122; Invitrogen) or mouse monoclonal anti-α-tubulin antibody (1:500 dilution: T9026; Sigma-Aldrich) in blocking buffer at 4°C overnight, and washed three times for 15 min in wash buffer (0.5% vol/vol Tween-20 in PBS). The gels were incubated with 8 μg/ml Atto 594 NHS-ester (Merck), 10 μg/ml Hoechst 33342 (Molecular Probes), and Alexa Fluor 488 goat anti-rabbit IgG (A11008; Invitrogen) or Alexa Fluor 488 goat anti-mouse IgG (A11001; Invitrogen) in PBS (1:500 dilution) at 37°C for 2.5 h followed by three washes of 15 min each in wash buffer (blocking and all antibody incubation steps were performed with gentle shaking). The gels were then washed three times for 15 min with wash buffer and expanded overnight in ultrapure water. The expanded gel was placed in a 35-mm glass-bottom dish (MatTek) with the 14-mm glass coated with poly-D-lysine. High-resolution images were acquired on a Zeiss Celldiscoverer 7 with Airyscan using a 50×/1.2 water objective. Confocal z-stacks were acquired using line scanning and the following settings: 55 × 55 nm pixel size, 170-nm z-step, 2.91 μs/pixel dwell time, 850 gain, and 3.5% (405 nm), 4.5% (488 nm), and 5.0% (561 nm) laser powers. The z-stack images were processed and analysed using Fiji (version 1.54f). Each experiment was repeated at least three times, and protein localisation was assessed in 5–10 cells per condition. Full imaging details are provided in the figure legends.

### Quantitative real-time PCR (qRT-PCR) analysis

Total RNA was extracted from WT and *Pbnek4-ko* zygotes at 2 hpa (three biological replicates per condition) using the RNA Purification Kit (Stratagene), and cDNA was synthesised using the RNA-to-cDNA Kit (Applied Biosystems). qPCR was performed with 80□ng of RNA using SYBR Green Fast Master Mix (Applied Biosystems) on an Applied Biosystems 7500 Fast system. Cycling conditions were: 95□°C for 20□s, followed by 40 cycles of 95□°C for 3□s and 60□°C for 30□s. Primers were designed using Primer3 (https://primer3.ut.ee/). Each gene was tested in three biological and technical replicates. *hsp70* (PBANKA_081890) and *arginyl-tRNA synthetase* (PBANKA_143420) were used as reference genes. Primer sequences are listed in Supplementary Table 17.

### Transcriptome study using RNA-Sequencing

RNA samples from WT and *Pbnek4-ko* zygotes at 2 hpa (three biological replicates per condition) were vacuum-concentrated by freeze drying and transported in RNA-stabilising tubes (Biomatrica) to ensure preservation. Strand-specific mRNA libraries were prepared using the TruSeq Stranded mRNA Library Prep Kit LT (Illumina). Paired-end sequencing (2 × 150 bp) was performed on an Illumina HiSeq 4000 platform. Quality control of raw sequencing reads was carried out using FASTQC (http://www.bioinformatics.babraham.ac.uk/projects/fastqc). Adapter trimming and removal of low-quality sequences were performed using Trimmomatic. Cleaned reads were aligned to the *P. berghei* ANKA reference genome (PlasmoDB release 40) using HISAT2 version 2.1.0, applying the parameter --rna-strandness FR. Gene-level expression was quantified using FeatureCounts. Raw read counts were converted to counts per million (CPM), and genes with CPM <1 in all but one replicate were excluded from downstream analysis. Library sizes were normalised using the trimmed mean of M-values (TMM) method in the EdgeR package. Processed count data were then analysed using the voom transformation from the limma package to model the mean–variance relationship. Differential expression analysis was performed with DESeq2, using a false discovery rate (FDR)-adjusted p-value threshold of 0.05 (Benjamini–Hochberg correction) and a fold-change cut-off of ≥2 to define significantly regulated genes.

### Immunoprecipitation and mass spectrometry

Purified gametocytes from WT-GFP and PbNEK4-GFP parasites were activated for 2 h and proteins were crosslinked using 1% formaldehyde for 10 min at room temperature. Excess aldehyde was quenched by incubating samples in 0.125 M glycine for 5 min, followed by three washes with PBS (pH 7.5). Immunoprecipitation of PbNEK4-GFP complexes was performed using the GFP-Trap_A kit (Chromotek) in accordance with the manufacturer’s protocol. GFP-bound protein complexes were captured on beads and processed for liquid chromatography–tandem mass spectrometry (LC-MS/MS). Beads were washed and incubated in ammonium bicarbonate (ABC) buffer. Reduction and alkylation were carried out using 10 mM TCEP (tris(2-carboxyethyl)phosphine hydrochloride) and 40 mM chloroacetamide (CAA) for 5 min at 70°C. Proteins were then digested with trypsin overnight at room temperature (1 μg trypsin per 100 μg of protein). Digestion was halted by adjusting the pH to 3–4 with 1% trifluoroacetic acid (TFA) before mass spectrometry analysis. Tryptic peptides were analysed via LC-MS/MS. Resulting raw data were searched using FragPipe version 21.0 (https://fragpipe.nesvilab.org/) against the PlasmoDB-66_PbergheiANKA_AnnotatedProteins database. Search parameters included trypsin digestion with up to two missed cleavages, fixed carbamidomethylation of cysteines, and variable modifications for methionine oxidation and N-terminal acetylation. Protein identification and quantification were performed using Scaffold (version 5.3.3, Proteome Software), and annotations were derived from PlasmoDB. Proteins were retained for analysis if they had ≥2 unique peptides and valid quantification in at least two replicates. The analysis was performed using two independent biological replicates.

### Phosphoproteomic analysis

#### Protein Extraction and Digestion via SDS-FASP

Zygotes of WT and *Pbnek4-ko* lines at 0 and 2 hpa were lysed in 400□μl of buffer containing 2% SDS, 25□mM NaCl, 50□mM Tris-HCl (pH□7.4), 2.5□mM EDTA, and 20□mM TCEP, supplemented with a protease and phosphatase inhibitor cocktail (Halt, Thermo Fisher Scientific). Samples were vortexed and heated at 95°C for 10 min with continuous mixing at 400□rpm. Genomic DNA was fragmented using four 10-second sonication pulses at 50% amplitude. Following centrifugation at 17,000□*g* for 30 min, supernatants were collected, and protein concentrations were determined using the Pierce 660nm Protein Assay Kit. For alkylation the volume corresponding to 1 mg of protein for each sample was adjusted to 300□μl and then incubated with 48□μl of 0.5□M iodoacetamide for 1 h at room temperature in the dark. Protein digestion was performed using the filter-aided sample preparation (FASP) method with Amicon Ultra-4 30□kDa centrifugal filter units (Millipore). Trypsin (Promega) was added at a 1:80 enzyme-to-protein ratio, and digestion proceeded overnight at room temperature. Resulting peptides were desalted using Pierce Peptide Desalting Spin Columns (Thermo Fisher Scientific) according to the manufacturer’s instructions and dried using a speed vacuum concentrator.

#### TMT-11plex Labelling

Peptide concentrations were measured using the Pierce Quantitative Colorimetric Peptide Assay (Thermo Fisher Scientific). For each sample, 160□μg of peptides were labelled with 800□μg of TMT reagents (Thermo Fisher Scientific), previously dissolved in 220□μl of 36% acetonitrile and 200□mM EPPS buffer (pH□8.5). Labelling reactions were carried out for 1 h at room temperature and quenched by adding hydroxylamine to a final concentration of 0.3% (v/v). Labelled samples were pooled, desalted using Pierce Peptide Desalting Spin Columns, and dried under vacuum.

#### Phosphopeptide Enrichment

Phosphopeptides were enriched using the High-Select Fe-NTA Phosphopeptide Enrichment Kit (Thermo Fisher Scientific) following the manufacturer’s protocol. Both the enriched phosphopeptide fractions and the flow-through were desalted using Pierce Peptide Desalting Spin Columns and dried using a speed vacuum concentrator.

#### LC-MS/MS Analysis

Dried peptides were reconstituted in 5% acetonitrile with 0.1% formic acid, and 2□μg of each sample was injected into a Vanquish NEO liquid chromatography system (Thermo Fisher Scientific) coupled to an Orbitrap Fusion Lumos Tribrid mass spectrometer (Thermo Fisher Scientific). Peptides were first trapped on an Acclaim PepMap 100 C18 trap column (75□μm × 20□mm, 3□μm particle size) and then separated on a homemade analytical column (75□μm × 500□mm) packed with ReproSil-Pur C18 resin (1.9□μm, 100□Å, Dr. Maisch GmbH). The analytical separation was run for 180 min using a gradient of H_2_O/FA 99.9%/0.1% (solvent A) and CH_3_CN/H_2_O/FA 80.0%/19.9%/0.1% (solvent B). The gradient was run from 5 % B to 28 % B in 160 min, then to 40% B in 20 min, then to 99% B in 10 min with a final stay of 20 min at 99 % B. Flow rate was of 250 nL/min an total run time was of 210 min. Data-Dependent Acquisition (DDA) was performed with MS1 full scan at a resolution of 120’000 FWHM followed by as many subsequent MS2 scans on selected precursors as possible within 3 second maximum cycle time. MS1 was performed in the Orbitrap with an AGC target of 4x10^5^, a maximum injection time of 50 ms and a scan range from 375 to 1500 m/z. MS2 was performed in the Orbitrap at a resolution of 50’000 FWHM using higher-energy collisional dissociation HCD at 38% NCE. Isolation windows was at 0.7 u with an AGC target of 5x10^4^ and a maximum injection time of 86 ms. A dynamic exclusion of parent ions of 60 s. with 10 ppm mass tolerance was applied.

#### Data Processing and Analysis

Raw data files were processed using Proteome Discoverer software version 2.4 (Thermo Fisher Scientific). Spectra were searched against the *P. berghei* ANKA protein database (https://PlasmoDB.org release 68), the *mus musculus* reference proteome database (UniProt, reviewed, release 2024_06, 17’207 entries) and an in-house database of common contaminants using the Mascot search engine (version 2.6.2, Matrix Science). Search parameters included trypsin as the digestion enzyme with up to one missed cleavage allowed, a precursor mass tolerance of 10□ppm, and a fragment mass tolerance of 0.02□Da. Carbamidomethylation of cysteine (+57.021□Da) and TMT modification of peptide N-termini and lysine residues (+229.163□Da) were set as fixed modifications, while oxidation of methionine (+15.995□Da) and phosphorylation of serine, threonine, and tyrosine (+79.966□Da) were set as variable modifications. Peptide-spectrum matches (PSMs) and peptides were validated using the Percolator algorithm, applying a false discovery rate (FDR) of 1%. Proteins were inferred from the identified peptides and filtered to achieve an FDR of 1%. Quantitative information was extracted using reporter ion intensities from TMT tags, and protein abundances were normalized based on the total peptide amount and scaled across all samples. Protein ratios were calculated using the summed abundances-based approach across groups, and statistical significance was assessed using ANOVA. For the analysis, three independent biological replicates were used for the 0 hpa *Pbnek4-ko*, 2 hpa WT, and 2 hpa *Pbnek4-ko* samples, while two independent biological replicates were used for the 0 hpa WT sample.

### Gene and protein annotation

Gene and protein annotations, including predicted functions and structural features, were primarily retrieved from PlasmoDB (release 68) and UniProt (version 23). Domain architectures and specific sequence motifs were predicted using InterPro (https://www.ebi.ac.uk/interpro/). For the uncharacterised conserved *Plasmodium* protein PBANKA_0806700, structural prediction was performed using the AlphaFold3 server (https://alphafoldserver.com/)^41^. The predicted structure was subsequently used as a query in Foldseek (https://search.foldseek.com/search)^42^ to identify structurally similar proteins, revealing the presence of putative HEAT-like repeats. Where applicable, functional annotations were based on previously published experimental characterisations or homology-based predictions, which are explicitly cited. A comprehensive summary of the key proteins highlighted in this study, including their identified domains, sizes, and the basis of their annotations, is provided in Supplementary Table 18.

## Data availability

The mass spectrometry proteomics data have been deposited to the ProteomeXchange Consortium via the PRIDE^84^ partner repository with the dataset identifier PXD070965 and 10.6019/PXD070965 (proteomics and phosphoproteomics) and PXD070161 and 10.6019/PXD070161 (GFP immunoprecipitations). RNAseq data have been deposited to the Gene Expression Omnibus under accession number the BioProject ID PRJNA1354107. The authors declare that all other relevant data generated or analysed during this study are included in the article or its supplementary information. Raw data are available for each figure. Materials are available from the corresponding authors on reasonable request.

## Extended Data Fig. legend

**Extended Data Fig. 1. *Plasmodium* NEK phylogeny, domain structure, function, and generation of PbNEK-GFP parasites. a.** Phylogenetic tree of NIMA-related protein kinases (NEKs) from *Plasmodium berghei* (Pb, light blue; PbNEK4, blue), *Plasmodium falciparum* (Pf, red), *Homo sapiens* (Hs, green), including NIMA (black) from *Aspergillus nidulans*. **b.** Domain structures of PbNEKs. **c.** Summary table of the function, expression stage, and localisation of PbNEKs. **d.** Schematic representation of the endogenous *nek4* locus, the GFP-tagging construct, and the recombined *nek4* locus following single homologous recombination. Arrows indicate the position of PCR primers used to confirm successful integration of the construct. **e.** Diagnostic PCR of *nek4* and WT-GFP parasites using the diagnostic primers to show the correct integration. Integration of the *nek4* tagging construct gives a band of ∼1400 bp. **f.** Western blot showing the expression of endogenous PbNEK4-GFP detected by anti-GFP antibody. The positions corresponding to the molecular weights of GFP (∼27 kDa) and PbNEK4-GFP (∼74 kDa) are indicated by arrows, respectively.

**Extended Data Fig. 2. PbNEK4-GFP localisation dynamics during zygote-ookinete development.** Live-cell imaging of PbNEK4-GFP location at different time points (minutes or hours post-activation, mpa or hpa). Representative live-cell images of different cells sampled at specific time points are shown. PbNEK4-GFP (green) parasites were labelled with Hoechst (blue) and a Cy3-conjugated 13.1 antibody (magenta), which recognises P28 protein on the surface of zygotes and ookinetes, immediately prior to imaging. Images are representative of 30–50 cells analysed across at least 3 independent biological experiments.

**Extended Data Fig. 3. Analyses of PbNEK4-GFP localisation dynamics. a.** Line graph showing the percentage of cells exhibiting a distinct PbNEK4-GFP focus in the nucleus (blue line) and at the cell surface (orange line) from pre-activation (0 hpa) to 24 hpa. Representative live-cell images of a 1 hpa zygote (PbNEK4-GFP alone and a GFP/Hoechst merge) are shown adjacent to the graph to illustrate the scoring. Orange and blue arrowheads indicate the distinct PbNEK4-GFP foci at the cell surface and within the nucleus, respectively. The y-axis represents the percentage of cells positive for the focal signal at each location. Scoring for the two locations was performed independently. Note that due to the spherical nature of the early zygote, simultaneously capturing both the nucleus and the cell surface in sharp microscopic focus (Z-plane) is technically challenging. Consequently, cells lacking an optimal focal plane for either region may have been scored as negative, meaning the percentages presented here might represent a slight underestimation. PbNEK4-GFP localisation was assessed in 30–50 individual cells per time point from at least three independent parasite preparation. **b.** Time-lapse imaging of a focal dot-like PbNEK4-GFP-positive structure in the Hoechst (blue)-stained PbNEK4-GFP (green) zygotes at 5 hpa. The focal dot-like PbNEK4-GFP-positive structure and the PbNEK4-GFP foci in the nucleus and at the cell periphery are indicated by yellow, magenta, and cyan arrowheads, respectively. Images are representative of 30–50 cells analysed across at least 3 independent biological experiments. **c.** Live-cell imaging of a focal dot-like PbNEK4-GFP-positive structure in the Hoechst (blue)-stained PbNEK4-GFP (green) developing ookinete at 10 hpa. The yellow arrowhead indicates the focal dot-like PbNEK4-GFP-posivite structure. Images are representative of 30–50 cells analysed across at least 3 independent biological experiments.

**Extended Data Fig. 4. Live-cell imaging and expansion microscopy of early zygotes. a.** PbNEK4-GFP and PbEB1-mCherry zygotes at 2 hpa. PbNEK4-GFP, PbEB1-mch, and colocalisation foci were indicated with green, magenta, and white arrow heads, respectively. Images are representative of at least 30 cells analysed across at least 3 independent biological experiments. **b.** Expansion microscopy of the zygote at 2 hpa displaying single MTOC with microtubule structures extending from it. Representative images from three independent biological experiments (n ≥ 5 cells examined). **c.** Zygotes activated for 2 hours displayed the development of the APC and MTOCs, along with microtubule structures extending form them. Representative images from three independent biological experiments (n ≥ 5 cells examined).

**Extended Data Fig. 5. Generation and genotypic and phenotypic analysis of *Pbnek4-ko* parasites. a.** Schematic representation of the endogenous *nek4* locus, the targeting knockout construct and the recombined *nek4* locus following double homologous crossover recombination. Arrows P1 and P2 indicate PCR primers used to confirm successful integration in the *nek4* locus following recombination, and arrows P3, 4, 5 and 6 indicate PCR primers used to show deletion of the *Pbnek4* gene. **b.** Integration PCR of the *nek4* locus in WT and *Pbnek4-ko* parasites using the primers, P1-6. **c.** TEM images of WT and *Pbnek4-ko* zygotes at 2 hpa. N, nucleus; APC, apical polar complex; A-Mt, APC microtubule; Pn-Mt, perinuclear microtubule; NM, nuclear membrane. Magenta box show condensed chromosomes. Representative images from three independent biological experiments (n ≥ 20 cells examined per condition). **d, e.** NHS-ester staining (gray) revealed that the apical polar complex (APC) had not yet formed in both WT (c) and *Pbnek4-ko* (d) female gametocytes before activation. A structure presumed to be the MTOC was also visible (magenta arrowhead), although it appeared smaller compared to that in zygotes at 2 hpa. Furthermore, anti α-tubulin antibody (green) and Hoechst (blue) staining revealed that no microtubule formation was observed around the potential MTOC and nucleus. Representative images from three independent biological experiments (n ≥ 5 cells examined per condition).

**Extended Data Fig. 6. qRT-PCR analysis of *Pbnek4-ko* and WT zygotes at 2 hpa and proteomic and phosphoproteomic analysis of *Pbnek4-ko* and WT zygotes at 0 hpa and *Pbnek4-ko* gametocytes at 0 hpa and *Pbnek4-ko* zygotes at 2 hpa. c.** Expression level validation of *nek4* and meiotic related genes using qRT-PCR in *Pbnek4-ko* and WT zygotes at 2 hpa. Shown is mean ± SEM; n = 3 independent experiments. Multiple comparisons *t* test, with post hoc test of Holm–Sidak showed significant differences in relative expression. *adjusted p-value□<□0.05, **adjusted p-value□<□0.01. **b-e.** Volcano plots displaying changes in total protein and phosphopeptide abundance between *Pbnek4-ko* and WT zygotes at 0 hpa and between *Pbnek4-ko* gametocytes at 0 hpa and *Pbnek4-ko* zygotes at 2 hpa. The log_2_ fold change (derived from the average of three biological replicates each) is plotted against the -log_10_ p-value. The dotted lines indicate the threshold for statistical significance (p-value ≤ 0.05 and fold change ≥ 1.5). Significantly upregulated and downregulated proteins/phosphopeptides are shown as red and blue dots, respectively. Proteins/phosphopeptides with non-significant changes are shown as grey dots. **b.** Comparison of protein abundance between *Pbnek4-ko* and WT zygotes at 0 hpa. **c.** Comparison of protein abundance between *Pbnek4-ko* gametocytes at 0 hpa and *Pbnek4-ko* zygotes at 2 hpa. **d.** Comparison of phosphopeptide abundance between *Pbnek4-ko* and WT zygotes at 0 hpa. **e.** Comparison of phosphopeptide abundance between *Pbnek4-ko* gametocytes at 0 hpa and *Pbnek4-ko* zygotes at 2 hpa. Key phosphopeptides from proteins of interest that are significantly upregulated in WT at 2 hpa (f) and downregulated in the *Pbnek4-ko* mutant at 2 hpa (g) are highlighted with yellow dots and protein names.

**Extended Data Fig. 7. PbDHC3-GFP localisation dynamics during zygote-ookinete development.** Live-cell imaging of PbDHC3-GFP location at different time points. Representative live-cell images of different cells sampled at specific time points are shown. PbDHC3-GFP (green) parasites were labelled with Hoechst (blue) and a Cy3-conjugated 13.1 antibody (magenta), which recognises P28 protein on the surface of zygotes and ookinetes, immediately prior to imaging. Magenta and cyan arrowheads indicate PbDHC3-GFP foci around the nucleus and PbDHC3-GFP localisation on the cortical region, respectively. Images are representative of 30–50 cells analysed across at least 3 independent biological experiments.

**Supplementary Table 1: RNA-Seq data analysis of *Pbnek4-ko* vs WT parasite 2 hpa**

**Supplementary Table 2: Gene Ontology Analysis of RNA-Seq data**

**Supplementary Table 3: Quantitative proteomics of WT 2 hpa versus WT 0 hpa parasites**

**Supplementary Table 4: Quantitative proteomics of *Pbnek4-ko* 2 hpa versus WT 2 hpa parasites**

**Supplementary Table 5: Quantitative proteomics of *Pbnek4-ko* 0 hpa versus WT 0 hpa parasites**

**Supplementary Table 6: Quantitative proteomics of *Pbnek4-ko* 2 hpa versus *Pbnek4-ko* 0 hpa parasites**

**Supplementary Table 7: Quantitative phosphoproteomics of WT 2 hpa versus WT 0 hpa parasites**

**Supplementary Table 8: Gene Ontology Analysis of phosphoproteomic data for WT 2 hpa versus WT 0 hpa parasites**

**Supplementary Table 9: Quantitative phosphoproteomics of *Pbnek4-ko* 2 hpa versus WT 2 hpa parasites**

**Supplementary Table 10: Gene Ontology Analysis of phosphoproteomic data for *Pbnek4-ko* 2 hpa versus WT 2 hpa parasites**

**Supplementary Table 11: Quantitative phosphoproteomics of *Pbnek4-ko* 2 hpa versus *Pbnek4-ko* 0 hpa parasites**

**Supp. Table 12: Quantitative phosphoproteomics of *Pbnek4-ko* 0 hpa versus WT 0 hpa parasites**

**Supplementary Table 13: GFP-Trap downs of PbNEK4-GFP parasites 2 hpa**

**Supplementary Table 14: Phosphatases and kinases identified in the *Pbnek4-ko* phosphoproteomics**

**Supplementary Table 15: AP2-Z target genes significantly altered in the *Pbnek4-ko* RNA-seq data**

**Supplementary Table 16: AP2-O target genes significantly altered in the *Pbnek4-ko* RNA-seq data**

**Supplementary Table 17: Primers used in this study**

**Supplementary Table 18: Summary of key proteins highlighted in this study and their functional annotations**

**Supplementary Video 1**

Time-lapse video microscopy of a focal dot-like PbNEK4-GFP-positive structure in the Hoechst (blue)-stained PbNEK4-GFP (green) zygotes at 5 hpa. Video is played back at 20× speed. Scale bar = 2 μm.

**Supplementary Video 2**

Time-lapse video microscopy of nuclear movement in the Hoechst (blue)-stained PbNEK4-GFP (green) zygotes at 2 hpa. Video is played back at 50× speed. Scale bar = 5 μm.

**Supplementary Video 3**

Time-lapse video microscopy of nuclear movement in the single zygote from Supplementary Video 1 (blue; Hoechst, green; PbNEK4-GFP). Video is played back at 50× speed. Scale bar = 5 μm.

**Supplementary Video 4**

Time-lapse video microscopy of nuclear movement in the Hoechst (blue)-stained PbEB1-GFP (green) zygotes at 2 hpa. Video is played back at 50× speed. Scale bar = 5 μm.

**Supplementary Video 5**

Time-lapse video microscopy of nuclear movement in the single zygote from Supplementary Video 3 (blue; Hoechst, green; PbEB1-GFP). Video is played back at 50× speed. Scale bar = 5 μm.

**Supplementary Video 6**

Time-lapse video microscopy of moving nuclei in the Hoechst (blue)-stained PbNEK4-GFP zygotes at 4 hpa. Video is played back at 20× speed. Scale bar = 5 μm.

**Supplementary Video 7**

Time-lapse video microscopy of static nuclei in the Hoechst (blue)-stained *Pbnek4-ko* zygotes at 4 hpa. Video is played back at 20× speed. Scale bar = 5 μm.

**Supplementary Video 8**

Time-lapse video microscopy of nuclear movement in the Hoechst (blue)-stained PbDHC3-GFP (green) zygotes at 3 hpa. Video is played back at 50× speed. Scale bar = 5 μm.

**Supplementary Video 9**

Time-lapse video microscopy of nuclear movement in the Hoechst (blue)-stained PbDHC3-GFP (green) zygotes at 5 hpa. Video is played back at 50× speed. Scale bar = 5 μm.

## Contributions

RT and DSG conceived the project. RY, RT, MZ, DB performed the live imaging, genotypic and phenotype analysis; RY, MH, DJF, SV performed TEM and SBFSEM analysis; SB,AN, KGLR performed RNA seq analysis; RY, DB, ARB and ECT performed proteomic analysis; CP, AH performed phosphoproteomics, RY, MZ performed U-ExM observation; RY, MH, MZ, DJPF, CP, AN, ECT, KGLR, AAH, SV, DSG, RT performed formal analysis; RT, DSG, KGLR, ECT acquired funding; RT, DSG and AAH supervised the project. RY, DSG, RT wrote the manuscript draft; All authors reviewed and edited the manuscript.

## Acknowledgements

RT is supported by an ERC advance grant funded by UKRI Frontier Science (EP/X024776/1), MRC UK (MR/K011782/1), and BBSRC (BB/L013827/1, BB/X014681/1). MZ, MH and DB were supported as research fellows and senior technician (EP/X024776/1). RY and DSG are supported by the BBSRC (BB/X014681/1). AAH is supported by the Francis Crick Institute (FC001097), which receives core funding from the Cancer Research UK (FC001097), the UK Medical Research Council (FC001097), and the Wellcome Trust (FC001097). KGLR is supported by the NIH/NIAID (R01 AI136511) and the University of California, Riverside NIFA-Hatch-225935. ECT was supported by a personal fellowship from the Nederlandse Organisatie voor Wetenschappelijk Onderzoek (NWO), the Netherlands (grant no. VI. Veni.202.223). We thank the School of Life Sciences Imaging (SLIM) for providing access to confocal and SIM microscopy and Dr Robert Markus for technical assistance. We thank the Nanoscale and Microscale Research Centre (nmRC) for providing access to instrumentation and Dr Michael W. Fey, Dr Julie Watts and Ms Nicola J. Weston for technical assistance. For Open Access, the authors have applied a CC BY public copyright licence to any Author Accepted Manuscript version arising from this submission. We thank Cleidiane Zampronio at Warwick University for mass spectrometry methods and Bio Support Unit, University of Nottingham for maintenance of mice used in this study.

## References

1. Bolcun-Filas E, Handel MA. Meiosis: the chromosomal foundation of reproduction. Biol Reprod 99, 112–126 (2018).

2. Marston AL, Amon A. Meiosis: cell-cycle controls shuffle and deal. Nat Rev Mol Cell Biol 5, 983–997 (2004).

3. Ohkura H. Meiosis: an overview of key differences from mitosis. Cold Spring Harb Perspect Biol 7, (2015).

4. Pesin JA, Orr-Weaver TL. Regulation of APC/C activators in mitosis and meiosis. Annu Rev Cell Dev Biol 24, 475–499 (2008).

5. Pereira C, Smolka MB, Weiss RS, Brieno-Enriquez MA. ATR signaling in mammalian meiosis: From upstream scaffolds to downstream signaling. Environ Mol Mutagen 61, 752–766 (2020).

6. Loidl J. Conservation and Variability of Meiosis Across the Eukaryotes. Annu Rev Genet 50, 293–316 (2016).

7. Mirzaghaderi G, Horandl E. The evolution of meiotic sex and its alternatives. Proc Biol Sci 283, (2016).

8. Guttery DS, Zeeshan M, Holder AA, Tromer EC, Tewari R. Meiosis in Plasmodium: how does it work? Trends Parasitol 39, 812–821 (2023).

9. Guttery DS, Zeeshan M, Ferguson DJP, Holder AA, Tewari R. Division and Transmission: Malaria Parasite Development in the Mosquito. Annu Rev Microbiol 76, 113–134 (2022).

10. WHO. World malaria report 2024. World Health Organisation **ISBN** 978-992-974-010444-010440, (2024).

11. Zickler D, Kleckner N. Meiosis: Dances Between Homologs. Annu Rev Genet 57, 1–63 (2023).

12. Sinden RE, Hartley RH, Winger L. The development of Plasmodium ookinetes in vitro: an ultrastructural study including a description of meiotic division. Parasitology 91 **(Pt** **2**), 227–244 (1985).

13. Sinden RE, Hartley RH. Identification of the meiotic division of malarial parasites. J Protozool 32, 742–744 (1985).

14. Sinden RE. Gametocytogenesis in Plasmodium spp., and observations on the meiotic division. Ann Soc Belg Med Trop 65 **Suppl 2**, 21–33 (1985).

15. Sinden RE, Canning EU, Spain B. Gametogenesis and fertilization in Plasmodium yoelii nigeriensis: a transmission electron microscope study. Proc R Soc Lond B Biol Sci 193, 55–76 (1976).

16. Zeeshan M, et al. Plasmodium ARK2 and EB1 drive unconventional spindle dynamics, during chromosome segregation in sexual transmission stages. Nat Commun 14, 5652 (2023).

17. Zeeshan M, et al. Real-time dynamics of Plasmodium NDC80 reveals unusual modes of chromosome segregation during parasite proliferation. J Cell Sci 134, (2020).

18. Guttery DS, et al. Genome-wide functional analysis of Plasmodium protein phosphatases reveals key regulators of parasite development and differentiation. Cell Host Microbe 16, 128–140 (2014).

19. Tewari R, et al. The systematic functional analysis of Plasmodium protein kinases identifies essential regulators of mosquito transmission. Cell Host Microbe 8, 377–387 (2010).

20. Ward P, Equinet L, Packer J, Doerig C. Protein kinases of the human malaria parasite Plasmodium falciparum: the kinome of a divergent eukaryote. BMC Genomics 5, 79 (2004).

21. Gao H, et al. ISP1-Anchored Polarization of GCbeta/CDC50A Complex Initiates Malaria Ookinete Gliding Motility. Curr Biol 28, 2763–2776 e2766 (2018).

22. Fry AM, Bayliss R, Roig J. Mitotic Regulation by NEK Kinase Networks. Front Cell Dev Biol 5, 102 (2017).

23. Fry AM, O’Regan L, Sabir SR, Bayliss R. Cell cycle regulation by the NEK family of protein kinases. J Cell Sci 125, 4423–4433 (2012).

24. Brieno-Enriquez MA, Moak SL, Holloway JK, Cohen PE. NIMA-related kinase 1 (NEK1) regulates meiosis I spindle assembly by altering the balance between alpha-Adducin and Myosin X. PLoS One 12, e0185780 (2017).

25. Reininger L, et al. A NIMA-related protein kinase is essential for completion of the sexual cycle of malaria parasites. J Biol Chem 280, 31957–31964 (2005).

26. Reininger L, Garcia M, Tomlins A, Muller S, Doerig C. The Plasmodium falciparum, Nima-related kinase Pfnek-4: a marker for asexual parasites committed to sexual differentiation. Malar J 11, 250 (2012).

27. Reininger L, et al. An essential role for the Plasmodium Nek-2 Nima-related protein kinase in the sexual development of malaria parasites. J Biol Chem 284, 20858–20868 (2009).

28. Zeeshan M, et al. Plasmodium NEK1 coordinates MTOC organisation and kinetochore attachment during rapid mitosis in male gamete formation. PLoS Biol 22, e3002802 (2024).

29. M’Saad O, Bewersdorf J. Light microscopy of proteins in their ultrastructural context. Nat Commun 11, 3850 (2020).

30. Cromer L, et al. Rapid meiotic prophase chromosome movements in Arabidopsis thaliana are linked to essential reorganization at the nuclear envelope. Nat Commun 15, 5964 (2024).

31. Christophorou N, Rubin T, Bonnet I, Piolot T, Arnaud M, Huynh JR. Microtubule-driven nuclear rotations promote meiotic chromosome dynamics. Nat Cell Biol 17, 1388–1400 (2015).

32. Saito TT, Tougan T, Okuzaki D, Kasama T, Nojima H. Mcp6, a meiosis-specific coiled-coil protein of Schizosaccharomyces pombe, localizes to the spindle pole body and is required for horsetail movement and recombination. J Cell Sci 118, 447–459 (2005).

33. Ding DQ, Chikashige Y, Haraguchi T, Hiraoka Y. Oscillatory nuclear movement in fission yeast meiotic prophase is driven by astral microtubules, as revealed by continuous observation of chromosomes and microtubules in living cells. J Cell Sci 111 **(Pt** **6**), 701–712 (1998).

34. Yamamoto A, West RR, McIntosh JR, Hiraoka Y. A cytoplasmic dynein heavy chain is required for oscillatory nuclear movement of meiotic prophase and efficient meiotic recombination in fission yeast. J Cell Biol 145, 1233–1249 (1999).

35. Vogel SK, Pavin N, Maghelli N, Julicher F, Tolic-Norrelykke IM. Self-organization of dynein motors generates meiotic nuclear oscillations. PLoS Biol 7, e1000087 (2009).

36. Yang S, et al. EB1 decoration of microtubule lattice facilitates spindle-kinetochore lateral attachment in Plasmodium male gametogenesis. Nat Commun 14, 2864 (2023).

37. Chikashige Y, et al. Telomere-led premeiotic chromosome movement in fission yeast. Science 264, 270–273 (1994).

38. Labella S, Woglar A, Jantsch V, Zetka M. Polo kinases establish links between meiotic chromosomes and cytoskeletal forces essential for homolog pairing. Dev Cell 21, 948–958 (2011).

39. Mlambo G, Coppens I, Kumar N. Aberrant sporogonic development of Dmc1 (a meiotic recombinase) deficient Plasmodium berghei parasites. PLoS One 7, e52480 (2012).

40. Tromer EC, Wemyss TA, Ludzia P, Waller RF, Akiyoshi B. Repurposing of synaptonemal complex proteins for kinetochores in Kinetoplastida. Open Biol 11, 210049 (2021).

41. Abramson J, et al. Accurate structure prediction of biomolecular interactions with AlphaFold 3. Nature 630, 493–500 (2024).

42. van Kempen M, et al. Fast and accurate protein structure search with Foldseek. Nat Biotechnol 42, 243–246 (2024).

43. Sayers C, et al. Systematic screens for fertility genes essential for malaria parasite transmission reveal conserved aspects of sex in a divergent eukaryote. Cell Syst 15, 1075–1091 e1076 (2024).

44. Hirai M, Maeta A, Mori T, Mita T. Pb103 Regulates Zygote/Ookinete Development in Plasmodium berghei via Double Zinc Finger Domains. Pathogens 10, (2021).

45. Nishi T, Kaneko I, Iwanaga S, Yuda M. Identification of a novel AP2 transcription factor in zygotes with an essential role in Plasmodium ookinete development. PLoS Pathog 18, e1010510 (2022).

46. Yuda M, Kaneko I, Murata Y, Iwanaga S, Nishi T. Mechanisms of triggering malaria gametocytogenesis by AP2-G. Parasitol Int 84, 102403 (2021).

47. Modrzynska K, et al. A Knockout Screen of ApiAP2 Genes Reveals Networks of Interacting Transcriptional Regulators Controlling the Plasmodium Life Cycle. Cell Host Microbe 21, 11–22 (2017).

48. Russell AJC, et al. Regulators of male and female sexual development are critical for the transmission of a malaria parasite. Cell Host Microbe 31, 305–319 e310 (2023).

49. Kaneko I, Nishi T, Iwanaga S, Yuda M. Differentiation of Plasmodium male gametocytes is initiated by the recruitment of a chromatin remodeler to a male-specific cis-element. Proc Natl Acad Sci U S A 120, e2303432120 (2023).

50. Hart KJ, Power BJ, Rios KT, Sebastian A, Lindner SE. The Plasmodium NOT1-G paralogue is an essential regulator of sexual stage maturation and parasite transmission. PLoS Biol 19, e3001434 (2021).

51. Drewes G, Ebneth A, Preuss U, Mandelkow EM, Mandelkow E. MARK, a novel family of protein kinases that phosphorylate microtubule-associated proteins and trigger microtubule disruption. Cell 89, 297–308 (1997).

52. Cao LS, Wang J, Chen Y, Deng H, Wang ZX, Wu JW. Structural basis for the regulation of maternal embryonic leucine zipper kinase. PLoS One 8, e70031 (2013).

53. Liu B, Liu C, Li Z, Liu W, Cui H, Yuan J. A subpellicular microtubule dynein transport machinery regulates ookinete morphogenesis for mosquito transmission of Plasmodium yoelii. Nat Commun 15, 8590 (2024).

54. Depoix D, et al. Vital role for Plasmodium berghei Kinesin8B in axoneme assembly during male gamete formation and mosquito transmission. Cell Microbiol 22, e13121 (2020).

55. van de Kooij B, et al. Comprehensive substrate specificity profiling of the human Nek kinome reveals unexpected signaling outputs. Elife 8, (2019).

56. Alexander J, et al. Spatial exclusivity combined with positive and negative selection of phosphorylation motifs is the basis for context-dependent mitotic signaling. Sci Signal 4, ra42 (2011).

57. Grallert A, Hagan IM. Schizosaccharomyces pombe NIMA-related kinase, Fin1, regulates spindle formation and an affinity of Polo for the SPB. EMBO J 21, 3096–3107 (2002).

58. Grallert A, Krapp A, Bagley S, Simanis V, Hagan IM. Recruitment of NIMA kinase shows that maturation of the S. pombe spindle-pole body occurs over consecutive cell cycles and reveals a role for NIMA in modulating SIN activity. Genes Dev 18, 1007–1021 (2004).

59. Yoshimura SH, Hirano T. HEAT repeats - versatile arrays of amphiphilic helices working in crowded environments? J Cell Sci 129, 3963–3970 (2016).

60. Ji DD, Sultan AA, Chakrabarti D, Horrocks P, Doerig C, Arnot DE. An RCC1-type guanidine exchange factor for the Ran G protein is found in the Plasmodium falciparum nucleus. Mol Biochem Parasitol 95, 165–170 (1998).

61. Zalli D, Bayliss R, Fry AM. The Nek8 protein kinase, mutated in the human cystic kidney disease nephronophthisis, is both activated and degraded during ciliogenesis. Hum Mol Genet 21, 1155–1171 (2012).

62. Burette M, et al. Two anchoring proteins control daughter apical complex assembly in Toxoplasma gondii. bioRxiv, (2026).

63. Koch LB, Spanos C, Kelly V, Ly T, Marston AL. Rewiring of the phosphoproteome executes two meiotic divisions in budding yeast. EMBO J 43, 1351–1383 (2024).

64. Sivakova B, et al. Quantitative proteomics and phosphoproteomics profiling of meiotic divisions in the fission yeast Schizosaccharomyces pombe. Sci Rep 14, 23105 (2024).

65. Swartz SZ, Nguyen HT, McEwan BC, Adamo ME, Cheeseman IM, Kettenbach AN. Selective dephosphorylation by PP2A-B55 directs the meiosis I-meiosis II transition in oocytes. Elife 10, (2021).

66. Okumura E, Morita A, Wakai M, Mochida S, Hara M, Kishimoto T. Cyclin B-Cdk1 inhibits protein phosphatase PP2A-B55 via a Greatwall kinase-independent mechanism. J Cell Biol 204, 881–889 (2014).

67. Kaneko I, Iwanaga S, Kato T, Kobayashi I, Yuda M. Genome-Wide Identification of the Target Genes of AP2-O, a Plasmodium AP2-Family Transcription Factor. PLoS Pathog 11, e1004905 (2015).

68. Murata Y, Nishi T, Kaneko I, Iwanaga S, Yuda M. Coordinated regulation of gene expression in Plasmodium female gametocytes by two transcription factors. Elife 12, (2024).

69. Anderson-White B, Beck JR, Chen CT, Meissner M, Bradley PJ, Gubbels MJ. Cytoskeleton assembly in Toxoplasma gondii cell division. Int Rev Cell Mol Biol 298, 1–31 (2012).

70. Tosetti N, et al. Essential function of the alveolin network in the subpellicular microtubules and conoid assembly in Toxoplasma gondii. Elife 9, (2020).

71. Harding CR, Frischknecht F. The Riveting Cellular Structures of Apicomplexan Parasites. Trends Parasitol 36, 979–991 (2020).

72. Burrell A, et al. Cellular electron tomography of the apical complex in the apicomplexan parasite Eimeria tenella shows a highly organised gateway for regulated secretion. PLoS Pathog 18, e1010666 (2022).

73. Chen CT, Gubbels MJ. TgCep250 is dynamically processed through the division cycle and is essential for structural integrity of the Toxoplasma centrosome. Mol Biol Cell 30, 1160–1169 (2019).

74. Hu X, O’Shaughnessy WJ, Beraki TG, Reese ML. Loss of the Conserved Alveolate Kinase MAPK2 Decouples Toxoplasma Cell Growth from Cell Division. mBio 11, (2020).

75. Guttery DS, et al. A unique protein phosphatase with kelch-like domains (PPKL) in Plasmodium modulates ookinete differentiation, motility and invasion. PLoS Pathog 8, e1002948 (2012).

76. Mytlis A, Levy K, Elkouby YM. The many faces of the bouquet centrosome MTOC in meiosis and germ cell development. Curr Opin Cell Biol 81, 102158 (2023).

77. Zeeshan M, et al. A novel SUN1-ALLAN complex coordinates segregation of the bipartite MTOC across the nuclear envelope during rapid closed mitosis in Plasmodium berghei. Elife 14, (2025).

78. Janse CJ, et al. High efficiency transfection of Plasmodium berghei facilitates novel selection procedures. Mol Biochem Parasitol 145, 60–70 (2006).

79. Winger LA, Tirawanchai N, Nicholas J, Carter HE, Smith JE, Sinden RE. Ookinete antigens of Plasmodium berghei. Appearance on the zygote surface of an Mr 21 kD determinant identified by transmission-blocking monoclonal antibodies. Parasite Immunol 10, 193–207 (1988).

80. Ferguson DJ, et al. Maternal inheritance and stage-specific variation of the apicoplast in Toxoplasma gondii during development in the intermediate and definitive host. Eukaryot Cell 4, 814–826 (2005).

81. Hair M, Moreira-Leite F, Ferguson DJP, Zeeshan M, Tewari R, Vaughan S. Atypical flagella assembly and haploid genome coiling during male gamete formation in Plasmodium. Nat Commun 14, 8263 (2023).

82. Yanase R, et al. Divergent Plasmodium kinases drive MTOC, kinetochore and axoneme organisation in male gametogenesis. Life Sci Alliance 8, (2025).

83. Liffner B, Silva T, Vega-Rodriguez J, Absalon S. Mosquito Tissue Ultrastructure-Expansion Microscopy (MoTissU-ExM) enables ultrastructural and anatomical analysis of malaria parasites and their mosquito. BMC Mehods 1, (2024).

84. Perez-Riverol Y, et al. The PRIDE database at 20 years: 2025 update. Nucleic Acids Res 53, D543–D553 (2025).

