## Extended Data Fig. 1-7 for "A divergent *Plasmodium* NEK4 acts as a key regulator driving the early events of meiosis"

**a**

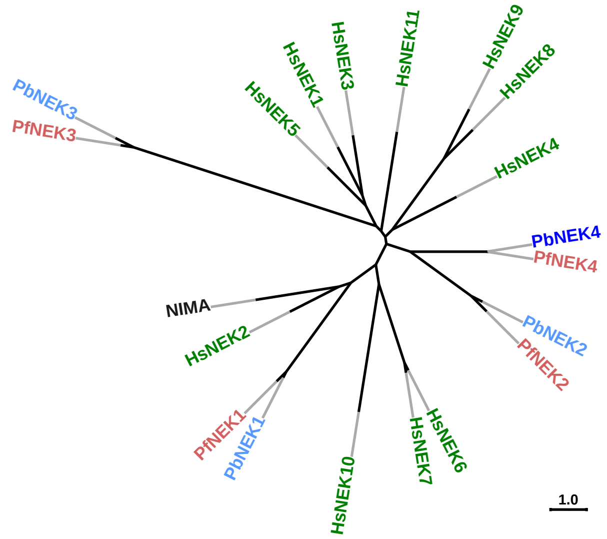


**b**

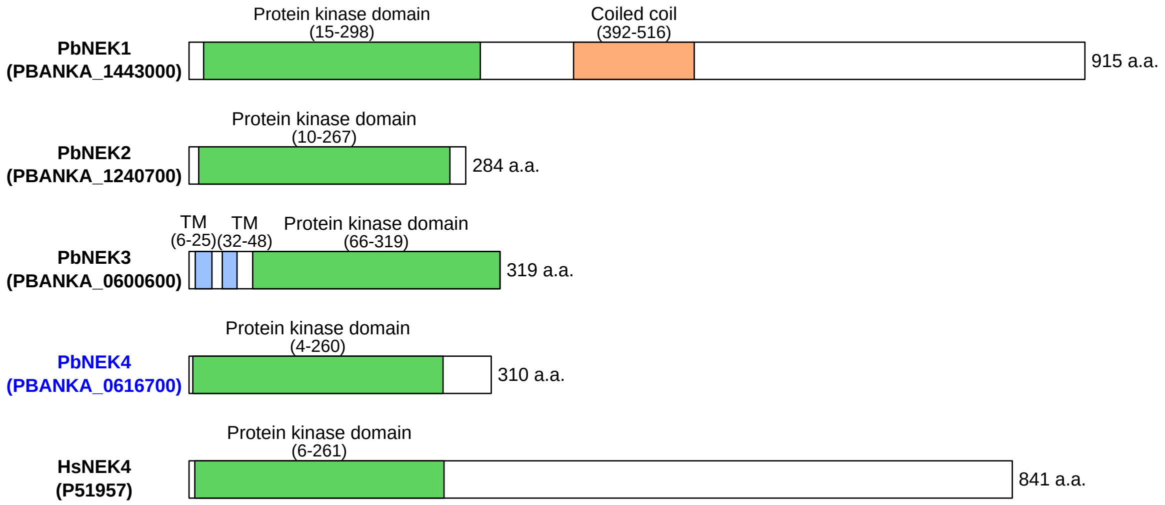


**c**

| Protein | Function | Cell stage/type | Localisation |
|---|---|---|---|
| PbNEK1 | Mitosis | Male gametocyte, blood stage cell | MTOC |
| PbNEK2 | Meiosis | Female gametocyte, zygote, ookinete | Subpellicular microtubule |
| PbNEK3 | Mitosis | Male gametocyte, blood stage cell | ? |
| PbNEK4 | Meiosis | Female gametocyte, zygote, ookinete | ? |

**d**

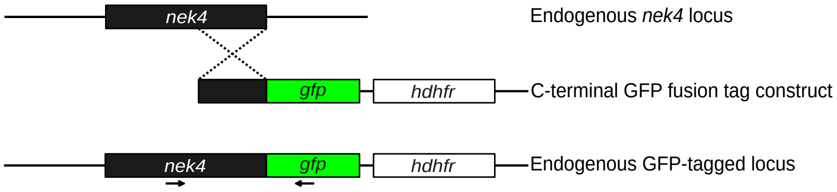


**e**

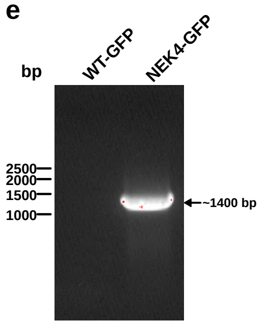


**f**

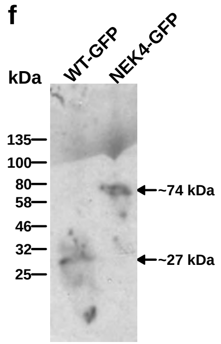

**Extended Data Fig. 2**

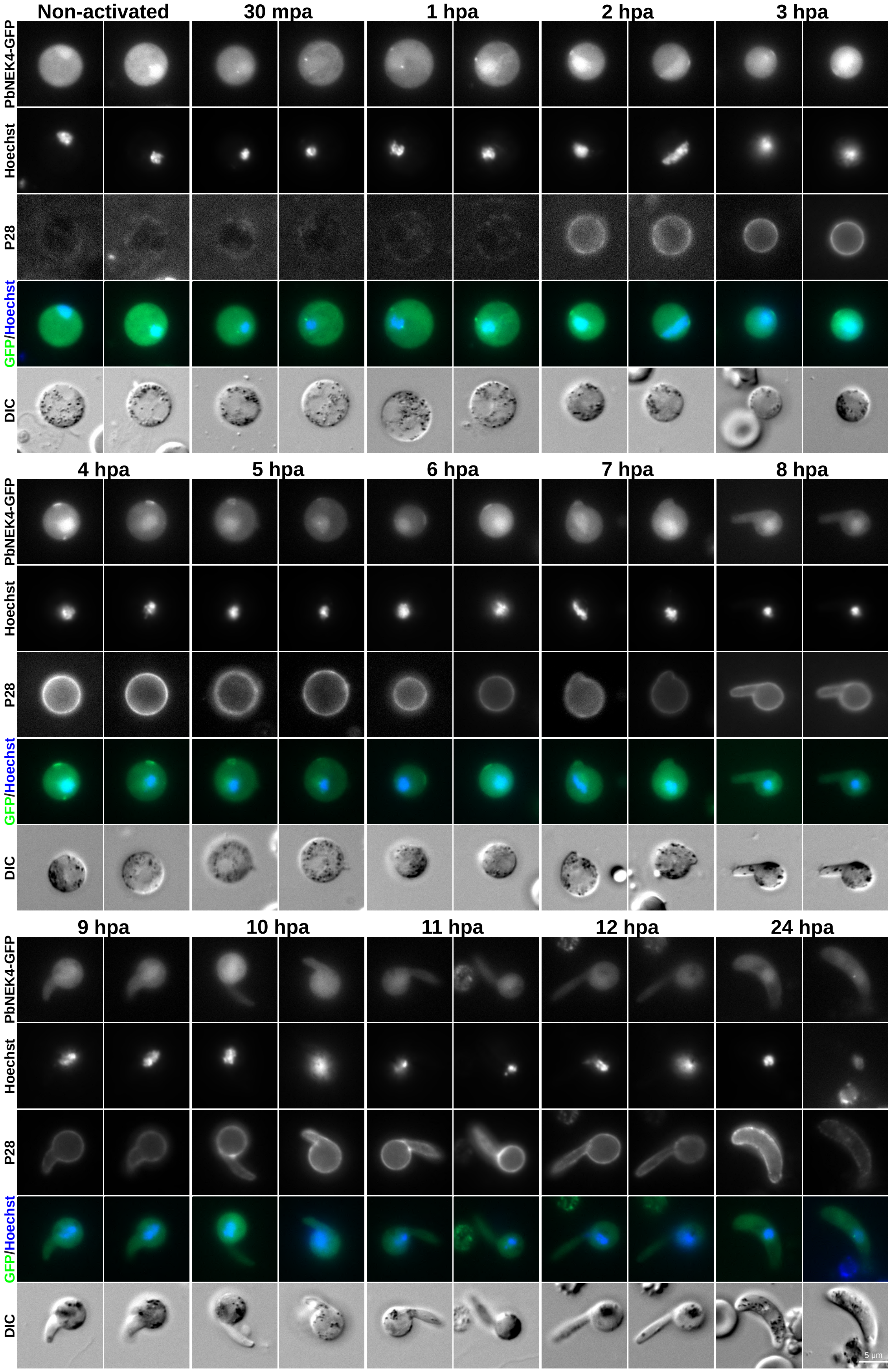

**Extended Data Fig. 3**

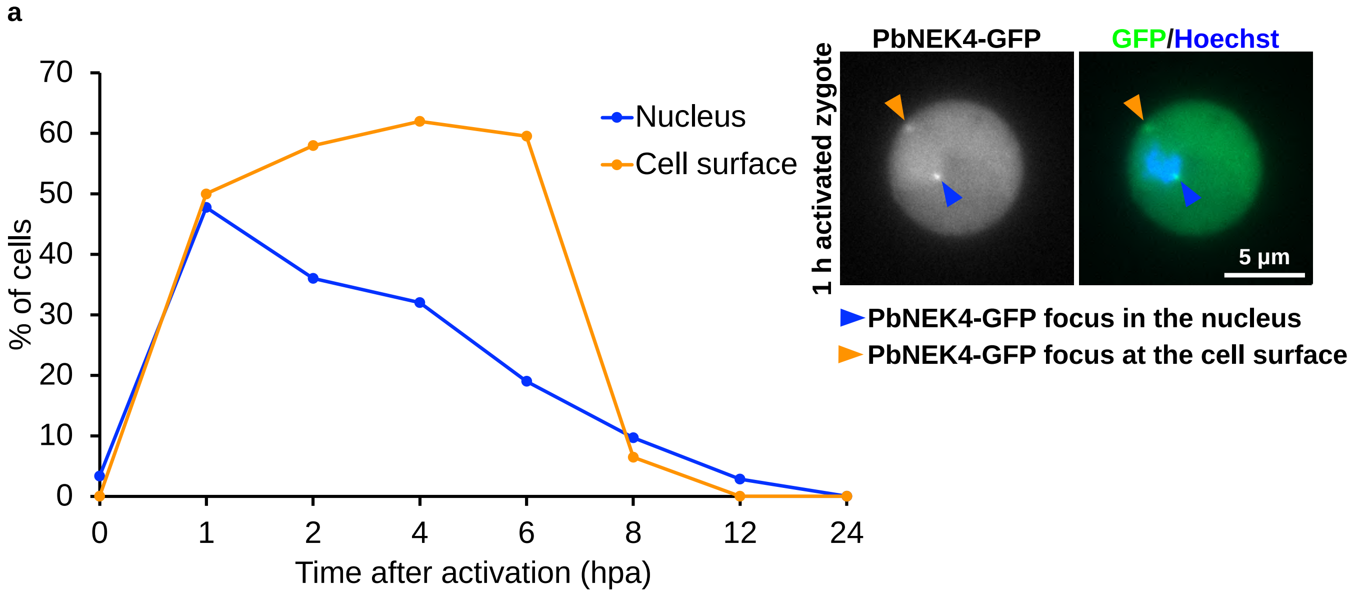


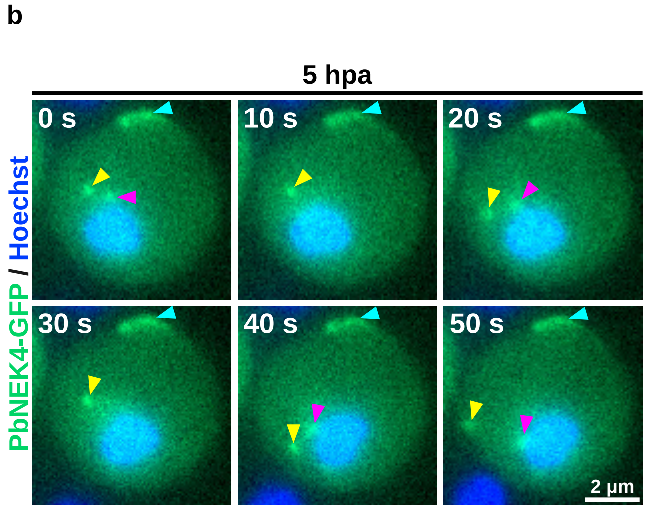


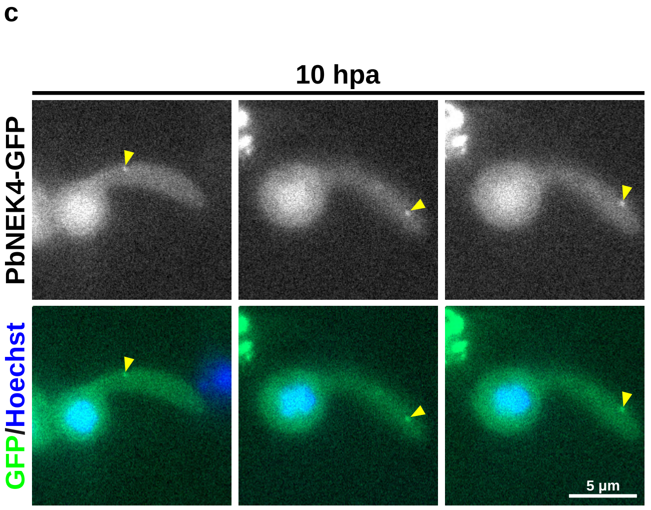

**Extended Data Fig. 4**

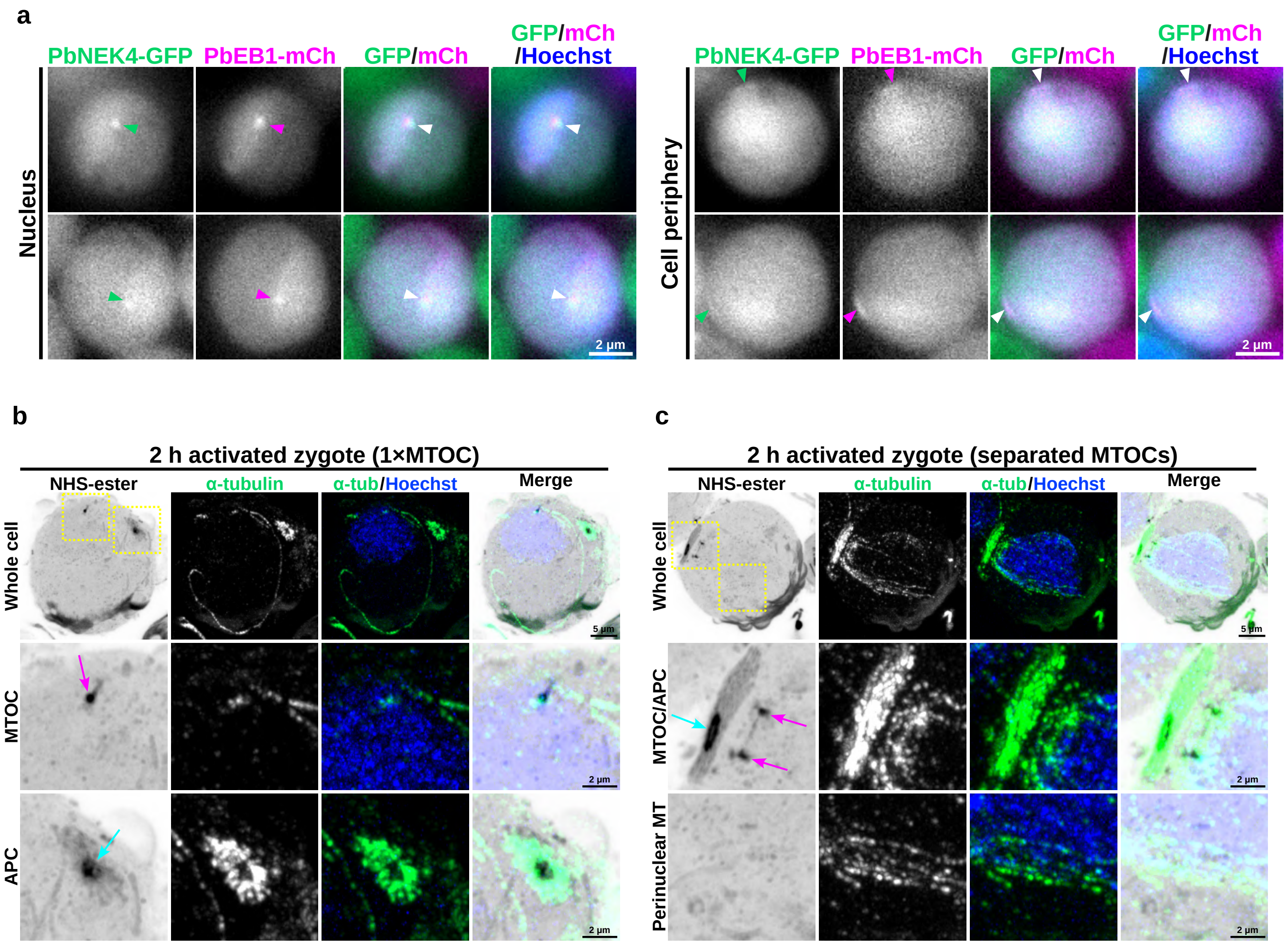

Extended Data Fig. 5

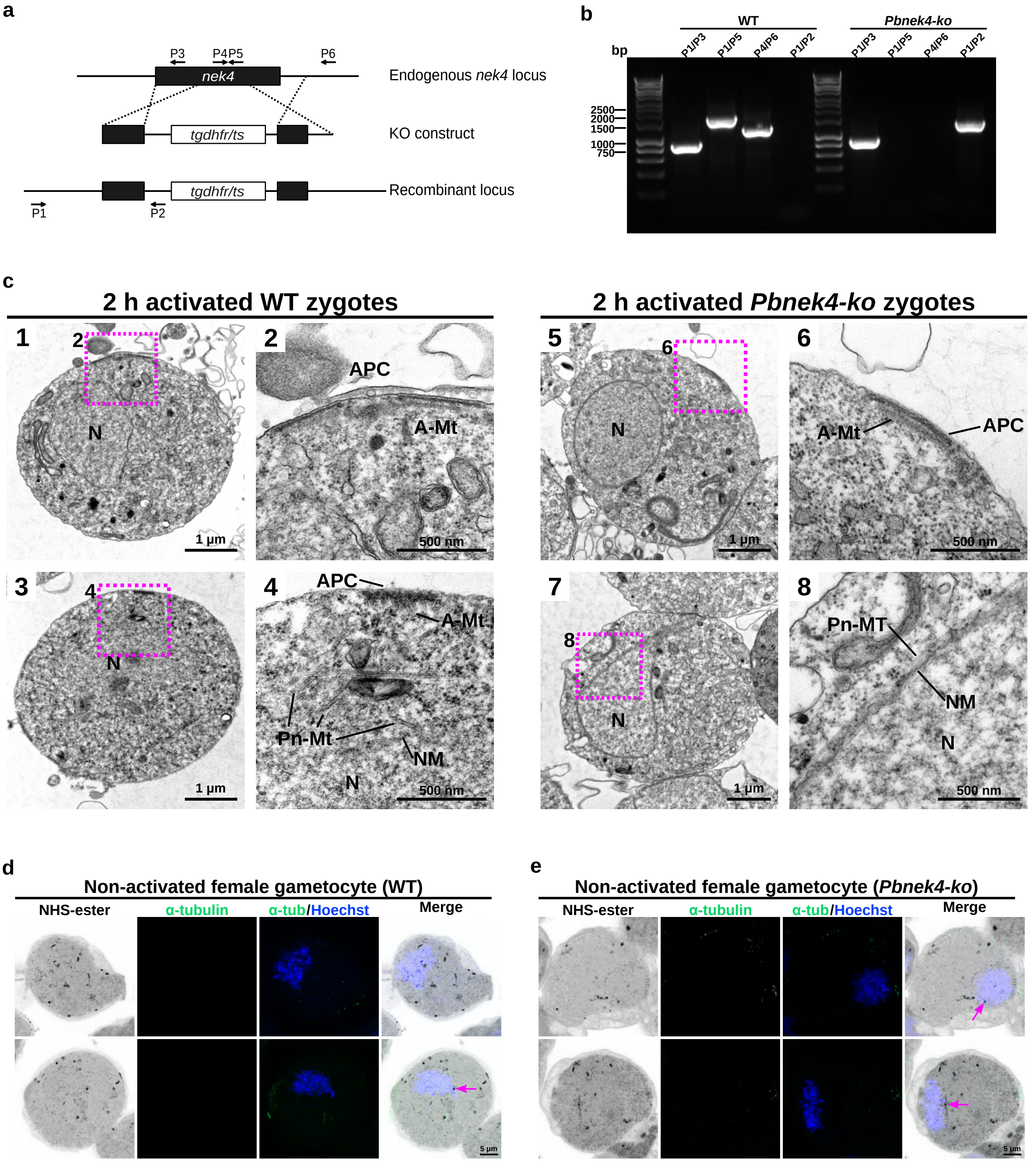

**Extended Data Fig. 6**

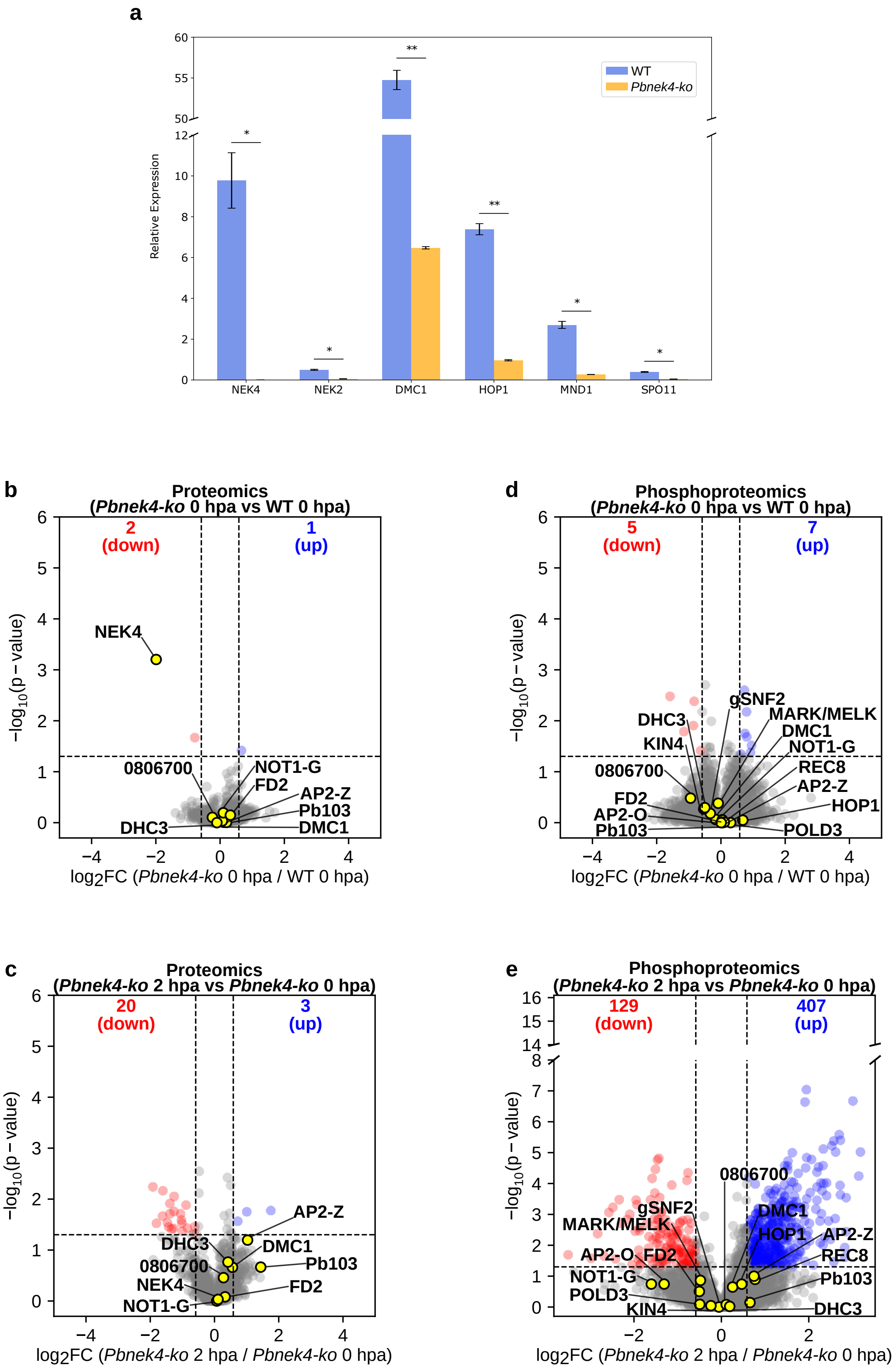

**Extended Data Fig. 7**

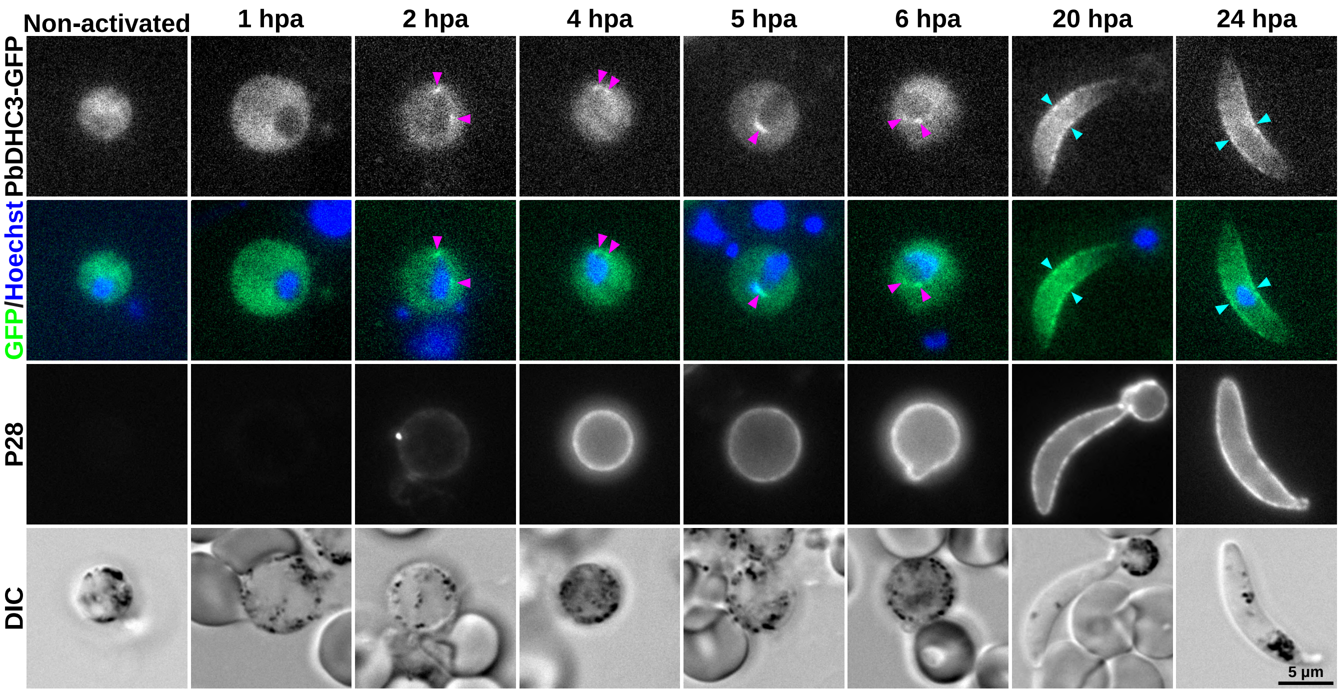
